# Proteasome-Bound UCH37 Debranches Ubiquitin Chains to Promote Degradation

**DOI:** 10.1101/2020.02.21.960088

**Authors:** Kirandeep K. Deol, Sean O. Crowe, Jiale Du, Heather Bisbee, Robert G. Guenette, Eric R. Strieter

**Author notes:** Correspondence: Eric R. Strieter.

## Abstract

The linkage, length, and architecture of ubiquitin (Ub) chains are all important variables in providing tight control over many biological paradigms. There are clear roles for branched architectures in regulating proteasome-mediated degradation, however the proteins that selectively recognize and process these atypical chains are unknown. Here, using synthetic and enzyme-derived ubiquitin chains along with intact mass spectrometry, we report that UCH37/UCHL5, a proteasome-associated deubiquitinase, exclusively cleaves K48 branched chains. The activity and selectivity toward branched chains is markedly enhanced by the proteasomal Ub receptor RPN13/ADRM1. Using proteasome complexes reconstituted with either active or inactive UCH37 together with protein substrates modified with branched chains, we find that chain debranching promotes degradation under multi-turnover conditions. These results are further supported by proteome-wide pulse-chase experiments, which show that the loss of UCH37 activity impairs global protein turnover. Our work therefore defines UCH37 as a debranching deubiquitinase important for promoting proteasomal degradation.

## INTRODUCTION

Ubiquitin (Ub) chains are a class of non-template-derived biopolymers with diverse structures that provide precise control over many biological pathways (Oh et al., 2018). The eight amino groups of Ub (M1, K6, K11, K27, K29, K33, K48, and K63), which decorate almost the entire surface, give rise to wide assortment of chain types (Swatek and Komander, 2016; Yau and Rape, 2016). Each one can be conjugated to the C-terminus of another Ub molecule to afford single-linkage (homotypic) and mixed-linkage (heterotypic) chains. Further expanding the diversity, heterotypic chains exist in different configurations: linear/unbranched and branched, with the latter composed of subunits that are modified at more than one amino group with other Ub molecules. Failure to build and remove specific chains at the appropriate time can be detrimental to cellular function (Damgaard et al., 2016; Heger et al., 2018; Popovic et al., 2014; Zhou et al., 2016), supporting the notion that the cellular fate of ubiquitinated proteins is largely dictated by chain type.

While efforts have largely focused on defining the functions of homotypic chains, heterotypic chains are emerging as important regulators of cellular pathways (Haakonsen and Rape, 2019). K11/K48 branched chains, for instance, increase in abundance during mitosis and proteotoxic stress to prioritize substrates, e.g., cell cycle regulators and misfolded proteins, for degradation by the 26S proteasome (Meyer and Rape, 2014; Samant et al., 2018; Yau et al., 2017). K29/K48 and K48/K63 branched chains have also been implicated in targeting proteins for degradation (Leto et al., 2019; Liu et al., 2017; Ohtake et al., 2018). M1/K63 chains, on the other hand, function independently of the proteasome (Emmerich et al., 2013; Wertz et al., 2015). Together, these results raise the possibility that branched conjugates can be selectively recognized by Ub-binding domains (UBDs) (Husnjak and Dikic, 2012) and processed by deubiquitinases (DUBs) (Mevissen and Komander, 2017). While there is evidence some UBDs may prefer branched conjugates (Boughton et al., 2020), a branched chain-selective DUB has not yet been identified.

UCH37/UCHL5 is a cysteine protease and member of the small family of DUBs referred to as the Ub C-terminal hydrolases (UCHs). Deficiencies in UCH37 lead to embryonic lethality in mice (Al-Shami et al., 2010) and overexpression has been found in several human cancers (Fang et al., 2012, 2013). At the cellular level, UCH37 has been implicated in TGF-β signaling (Wicks et al., 2005, 2006), Wnt signaling (Han et al., 2017), DNA double-strand break repair (Nishi et al., 2014), cell cycle progression (Randles et al., 2016), NF-κB activation (Mazumdar et al., 2010), and adipogenesis (van Beekum et al., 2012). Most of these functions have been attributed to UCH37’s association with either the proteasome (Hamazaki et al., 2006; Jørgensen et al., 2006; Qiu et al., 2006; Yao et al., 2006) or INO80 chromatin remodeling complex (Yao et al., 2008). In the case of the proteasome, the dogma is that UCH37 trims homotypic K48 chains to rescue proteins from degradation (Lam et al., 1997). The kinetics of homotypic K48 chain cleavage, however, are exceedingly slow (Bett et al., 2015; Lu et al., 2017; Yao et al., 2006). Moreover, purified human proteasomes with UCH37 as the only non-essential DUB are unable to cleave K48 chains (Lu et al., 2015b). Thus, the function of UCH37 on the proteasome remains poorly understood.

In the present study, we demonstrate that UCH37 exclusively cleaves branched Ub chains. Using a library of designer Ub chains, we show that K48 linkages are readily removed when presented in the context of a branched chain. Middle-down mass spectrometry combined with quantitative linkage analysis reveals that the K48-specific debranching activity extends to complex chain mixtures. The proteasomal subunit RPN13/ADRM1 markedly enhances both the activity and selectivity of UCH37. In the context of the proteasome, debranching activity is not only retained but also important for the degradation of substrate proteins modified with K48-containing branched chains. Proteome-wide analysis of global protein turnover further substantiates these findings by demonstrating that proteins are turned over more efficiently in cells expressing catalytically active UCH37 compared to cells with inactive UCH37. Our work therefore identifies UCH37 as a chain debranching enzyme and uncovers the mechanism by which UCH37 regulates proteasomal degradation.

## RESULTS

### UCH37 Cleaves K48 Branched Ub Trimers

Our lab developed a straightforward method for synthesizing a diverse set of Ub homo-oligomers based on thiol-ene coupling (TEC) (Figure 1A) (Trang et al., 2012; Valkevich et al., 2012). Utilizing this library of TEC-derived Ub homo-oligomers, we identified branched conjugates bearing K48 linkages as potential targets of UCH37 (Figure 1B). To confirm this result, a branched trimer bearing native isopeptide bonds was synthesized and subjected to UCH37. We found that UCH37 cleaves the native branched trimer in a time- and concentration-dependent manner (Figures 1C and D), suggesting branched Ub homo-oligomers could be the principal target of UCH37.

**Figure 1.**
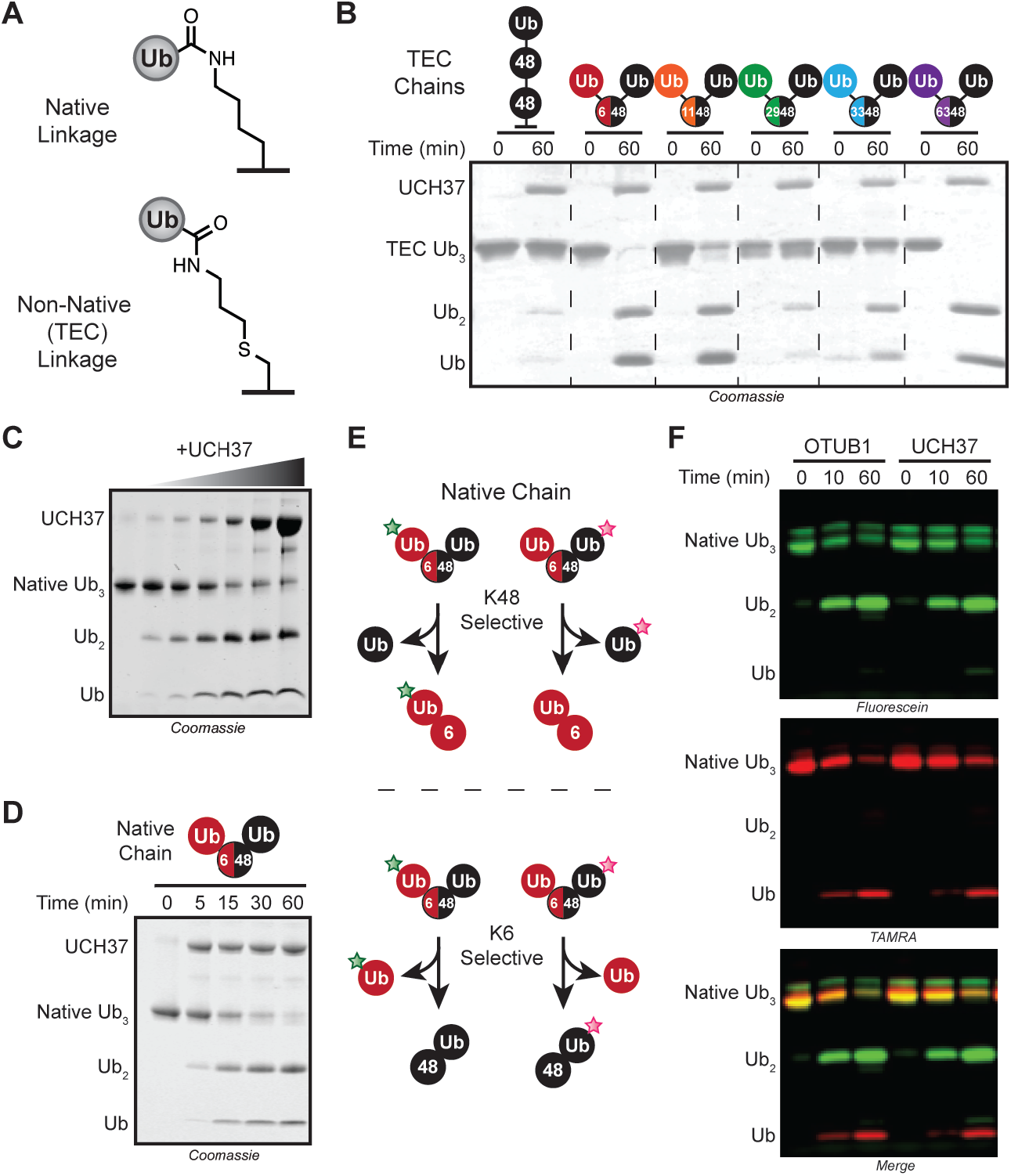
UCH37 Cleaves K48 Linkages in Branched Ubiquitin Trimers. (A) Structures of both native and thiol-ene coupled (TEC) isopeptide bonds used in this study. (B) SDS-PAGE analysis of TEC-derived branch tri-Ub (10 μM) with UCH37 (1 μM). Linkages in branched tri-Ub are represented by X/Y, where X and Y denote the positions of the isopeptide bonds. (C) SDS-PAGE analysis of native K6/K48 branch tri-Ub (10 μM) with varying concentrations of UCH37 (0, 0.1, 0.25, 0.5, 1, 5, and 10 μM). (D) SDS-PAGE analysis of the time course for the UCH37-catalyzed cleavage of native K6/K48 branch tri-Ub (10 μM). (E) Schematic for subunit-specific labeling of native K6/K48 branch tri-Ub with different fluorophores to report on the linkage specificity of UCH37. (F) Fluorescence analysis of cleavage reactions with either OTUB1 (5 μM) or UCH37 (5 μM) and fluorophore-labeled native K6/K48 branch tri-Ub (10 μM). See also Figure S1.

To further explore this unique reactivity, we wanted to determine whether UCH37 also exhibits linkage specificity. When UCH37 encounters a branch point, there are two possible linkages that can be cleaved. We used a sortagging approach to selectively label individual Ub subunits of a native chain with different fluorophores (Crowe et al., 2016). With a fluorescent native K6/K48 branched trimer as a substrate, the logic was that K48 cleavage should furnish a fluorescein-labeled di-Ub and TAMRA-labeled mono-Ub (Figures 1E). By contrast, scission of the K6 linkage should yield a TAMRA-labeled di-Ub and fluorescein-labeled mono-Ub, thus allowing us to interrogate UCH37 linkage selectivity. The results are consistent with K48 linkage specificity, as the products from UCH37-mediated cleavage mirror those generated by the K48 linkage specific DUB OTUB1 (Figure 1F). Cleavage patterns of TEC-derived chains also support this conclusion (Figures S1A-B). Replacing the non-K48 distal subunit with either GFP or SUMO abrogates cleavage, indicating that all three subunits in the chain must be Ub (Figure S1D). Together, our results show that UCH37 has the unprecedented ability to exclusively target K48 branch points for cleavage.

### UCH37 Removes K48 Branch Points in Complex Mixtures of Chains

Although Ub trimers are good model systems, they do not reflect the heterogeneity of ubiquitination observed in cells or in vitro. Thus, we sought to analyze the debranching activity of UCH37 in the context of heterogeneous chain populations where there is considerable variability in chain length and frequency of branch points. Identifying branch points in complex mixtures of chains, however, is challenging. Chain restriction analysis does not inform on architecture and multiple modifications on a single polypeptide chain are difficult to detect using standard bottom-up proteomic approaches (Mevissen et al., 2013). We turned to intact mass spectrometry (MS), as this has proven to be a powerful method for identifying and characterizing branched conjugates (Swatek et al., 2019; Valkevich et al., 2014).

A series of enzymatic reactions were performed based on their ability to generate chains with a specific mixture of linkages (Figure 2A). In each case, high molecular weight (HMW) conjugates were isolated and analyzed by intact MS. The extent of branching (2xdiGly-Ub_1-74_) varies from 4-14% of the total Ub population (Figures 2B-D; top spectra). According to electron capture dissociation (ECD) and electron transfer dissociation (ETD) analysis of the 2xdiGly-Ub_1-74_ peak, NleL generates K6/K48 branch points (Hospenthal et al., 2013), the combination of UBE2S (Bremm et al., 2010) and UBE2R1 forms K11/K48 branches, and UBE2N/UBE2V2 with UBE2R1 builds K48/K63 bifurcations (Nakasone et al., 2013) (Figures S2B-D).

**Figure 2.**
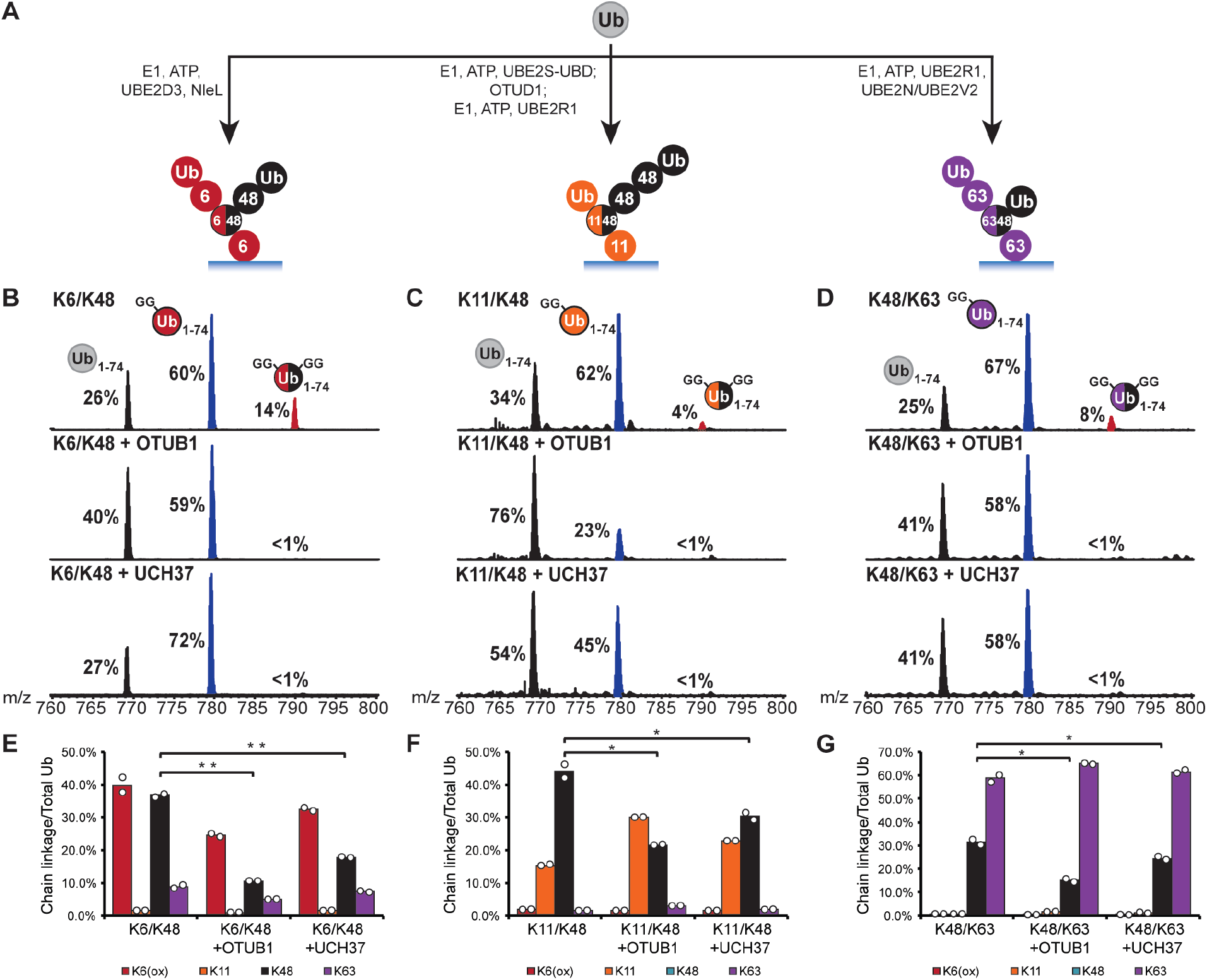
UCH37 Removes K48 Branch Points in Complex Chain Mixtures. (A) Schematic showing the assembly of high molecular weight ubiquitin chain mixtures used in this study. (B-D) Ub middle-down (Ub MiD) MS analysis of HMW K6/K48, K11/K48, and K48/K63 chains (top) treated with either OTUB1 (3 μM, middle) or UCH37 (3 μM, bottom). Percentages correspond to the relative quantification values of the 11+ charge state for each Ub species: Ub_1-74_, 1xdiGly-Ub_1-74_, and 2xdiGly-Ub_1-74_. (E-G) Ub-AQUA analysis of HMW K6/K48, K11/K48, and K48/K63 chains before and after OTUB1 (15 μM, middle) or UCH37 (3 μM, last) treatment. For all points, \**P*<0.025, \*\**P*<0.01 (Student’s T-test) All MS spectra are representative traces and quantification values (E-G) are derived from averaging fits of 2 independent experiments shown with SEM. See also Figure S2 and Table S1A-C.

MS analysis of the HMW conjugates after the addition of UCH37 shows a complete loss of the 2xdiGly-Ub_1-74_ species indicative of debranching (Figures 2B-D; bottom spectra). With K6/K48 chains, the disappearance of branched chains (2xdiGly-Ub_1-74_) coincides with an increase in the linear/unbranched chains (diGly-Ub_1-74_) (Figure 2B; bottom spectrum). Subjecting the same chains to OTUB1, which should be immune to chain architecture and cleave any K48 linkage, decreases the branch point and increases the relative amount of mono-Ub/end caps (Ub_1-74_) (Figure 2B; middle spectrum). With K11/K48 and K48/K63 chains, both UCH37-catalyzed debranching and the global removal of K48 linkages by OTUB1 afford an increase in Ub_1-74_ (Figures 2C-D; middle and bottom spectra). Chains built with non-K48 linkages, e.g., K11 and K63, are not targeted for cleavage by UCH37 (Figures S2E-F).

To determine whether UCH37 retains specificity toward K48 linkages in these complex chains, we used isotopically labeled, absolute quantitation (AQUA) peptide standards for each linkage of Ub (Kirkpatrick et al., 2006). HMW conjugates were analyzed both prior to and after the addition of OTUB1 and UCH37. The most significant changes occur in the K48 levels for each set of chains, indicating this linkage is the primary target of UCH37 (Figures 2E-G). The K48-specific debranching activity observed with model Ub trimers thus holds true for more complex chains.

### RPN13 Enhances the Debranching Activity of UCH37

Our data with free UCH37 suggest it could function as a chain debranching enzyme; however, there is little evidence that UCH37 acts on its own. One of its primary binding partners is the proteasomal Ub receptor RPN13 (Figure 3A) (Hamazaki et al., 2006; Jørgensen et al., 2006; Qiu et al., 2006; Yao et al., 2006). Previous studies have shown that RPN13 enhances UCH37’s ability to cleave the fluorogenic substrate Ub-AMC (Sahtoe et al., 2015; VanderLinden et al., 2015). Thus, we wanted to evaluate the effects of RPN13 on chain debranching. We first attempted to reconstitute the UCH37•RPN13 complex by mixing the purified recombinant proteins; however, we did not observe activity. We then co-purified UCH37 and full length RPN13 (Figure S3A). The resulting UCH37•RPN13 complex exhibits higher activity relative to free UCH37 against Ub-AMC, branched trimers, and HMW chains (Figures 3B, 3D and S3B). Analysis of cleavage reactions with HMW chains confirmed that the UCH37•RPN13 complex displays the same reactivity toward K48 branch points as free UCH37 (Figures 3C and S3C-D).

**Figure 3.**
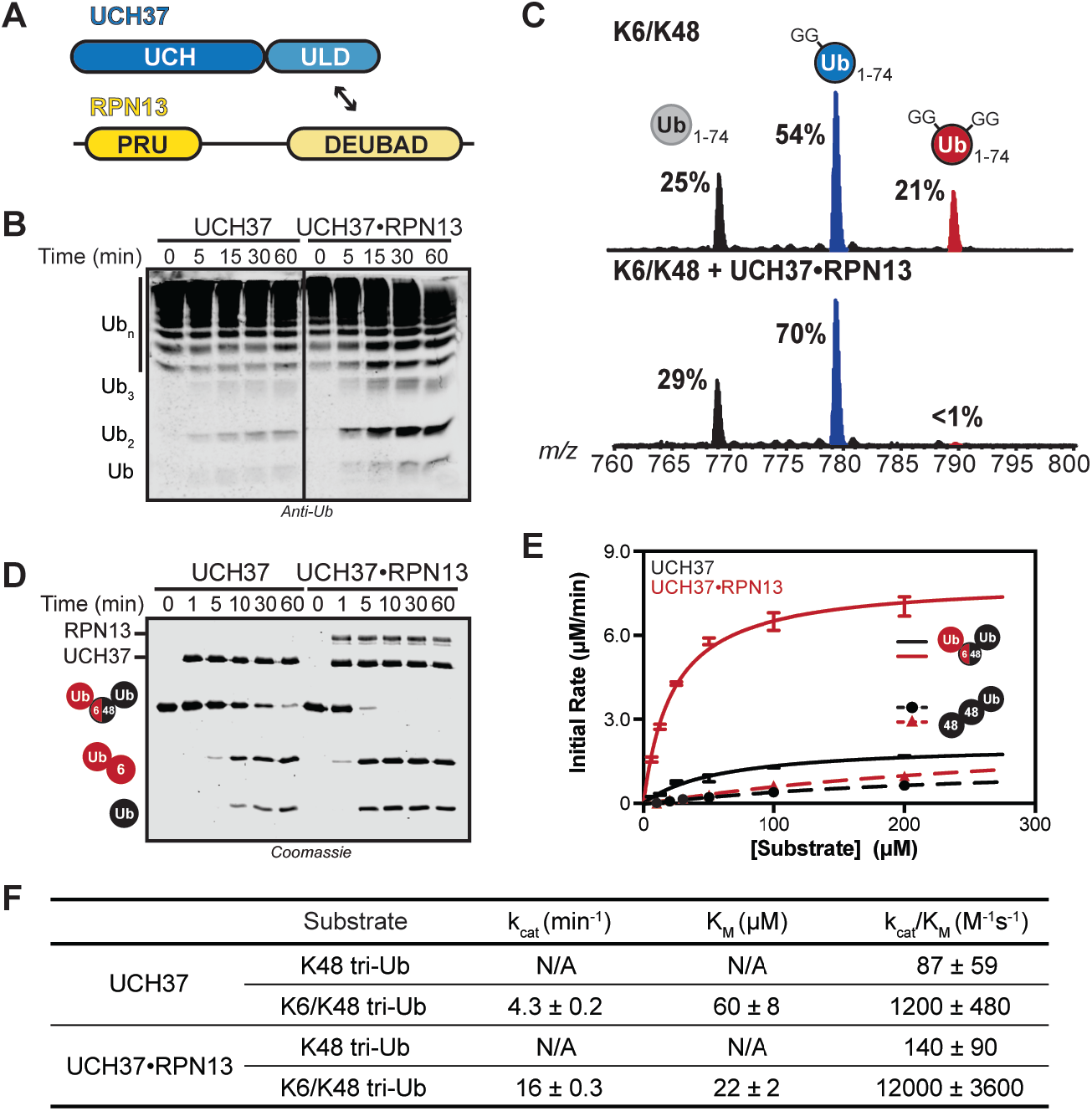
Steady-State Kinetic Analysis of Chain Debranching. (A) Constructs used in this study. (B) Western blot analysis of cleavage reactions with HMW K6/K48 chains. Reactions are performed with either UCH37 (1 µM) or UCH37•RPN13 complex (1 µM), tracked by SDS-PAGE, and visualized using the α-Ub P4D1 antibody. (C) Ub MiD MS analysis of HMW K6/K48 chains (top) followed by treatment with UCH37•RPN13 (1 μM, bottom). Percentages correspond to the relative quantification values of the 11+ charge state for each Ub species: Ub_1-74_, 1xdiGly-Ub_1-74_, and 2xdiGly-Ub_1-74_. (D) SDS-PAGE analysis of K6/K48 branched tri-Ub (10 μM) hydrolysis by UCH37 (1 μM) or UCH37•RPN13 (1 μM). (E) Michaelis-Menten plot for the hydrolysis of native K6/K48 branched tri-Ub (solid line) or K48 tri-Ub (dashed line) by either free UCH37 (black) or UCH37•RPN13 (red). Enzyme concentrations are 0.5 μM for native K6/K48 branched tri-Ub and 1 μM for K48 tri-Ub. (F) Table of kinetic parameters measured for all experiments following the initial rates of di-Ub formation. All kinetic curves are averaged representative traces and constants (E-F) are derived from averaging fits of independent experiments with SD (n = 3). See also Figure S3.

Encouraged by these results, we sought to obtain more quantitative information. With the native K6/K48 branched trimer as a model substrate we measured the initial rates of debranching using a gel-based assay. The formation of both di-Ub and mono-Ub were monitored and the resulting data were fit to the Michaelis-Menten equation (Figure 3E). The steady-state parameters show that the presence of RPN13 confers a 2-fold decrease in *K*_m_ and a 6-fold increase in *k*_cat_ (Figure 3F). RPN13 therefore boosts the debranching activity of UCH37 by an order of magnitude from 1200 M^−1^•s^−1^ to 12000 M^−1^•s^−1^. Interestingly, RPN13 has little effect on the cleavage efficiency of homotypic K48 chains. In fact, comparing *k*_cat_/*K*_m_ for the cleavage of K6/K48 tri-Ub and K48 tri-Ub reveals that the selectivity for the branched trimer increases from 12- to 90-fold in the presence of RPN13.

The heightened activity bestowed on UCH37 by RPN13 can largely be ascribed to the DEUBAD (DEUBiquitinase ADaptor) domain. The C-terminal DEUBAD domain is conserved among other UCH regulatory proteins (Sanchez-pulido et al., 2012) and is necessary and sufficient for enhancing UCH37-catalyzed hydrolysis of an artificial Ub-AMC substrate (Sahtoe et al., 2015; VanderLinden et al., 2015). Steady-state kinetic analysis of the debranching reaction with UCH37 and DEUBAD alone reveals a slight decrease in *k*_cat_ relative to full-length RPN13 (∼1.6-fold), but the *K*_m_ remains relatively the same (Figure S3C). Similar kinetics are observed when UCH37 is bound to a mutant form of RPN13 that is incapable of binding Ub (RPN13^L56A/F76R/D79N^; Lu et al., 2017; Schreiner et al., 2008) (Figure S3C). These results suggest that Ub-binding by the PRU (Pleckstrin-like Receptor for Ub) domain of RPN13 could contribute some degree to the catalytic events following Michaelis complex formation.

### Homotypic K48 Chains Inhibit Debranching *In Trans* but not *In Cis*

That UCH37•RPN13 cleaves homotypic K48 chains at all suggests that unbranched chains might interfere with its debranching activity. K48 tetra-Ub chains were added *in trans* at varying concentrations to reactions with K6/K48 tri-Ub as the substrate (Figure 4A). Global fitting of the steady-state kinetic data shows that K48 tetra-Ub acts as a competitive inhibitor (Figures 4B and S4A). The *K*_i_ of 0.7 μM for K48 tetra-Ub is approximately 31-fold lower than the *K*_m_ for K6/K48 tri-Ub with no inhibitor. Analysis of binding by isothermal titration calorimetry (ITC) confirms that the active UCH37•RPN13 complex binds K48 tri- and tetra-Ub with rather high affinity (*K*_d_ = 0.33 μM for tri-Ub and 0.15 μM for tetra-Ub) (Figures 4C-D). Similar binding strengths are observed for the inactive UCH37 C88A•RPN13 complex, as evidenced by both ITC (Figures S4B-E) and fluorescence polarization (Du and Strieter, 2018). Together, these results argue that homotypic K48 chains are potent inhibitors of debranching *in trans*.

**Figure 4.**
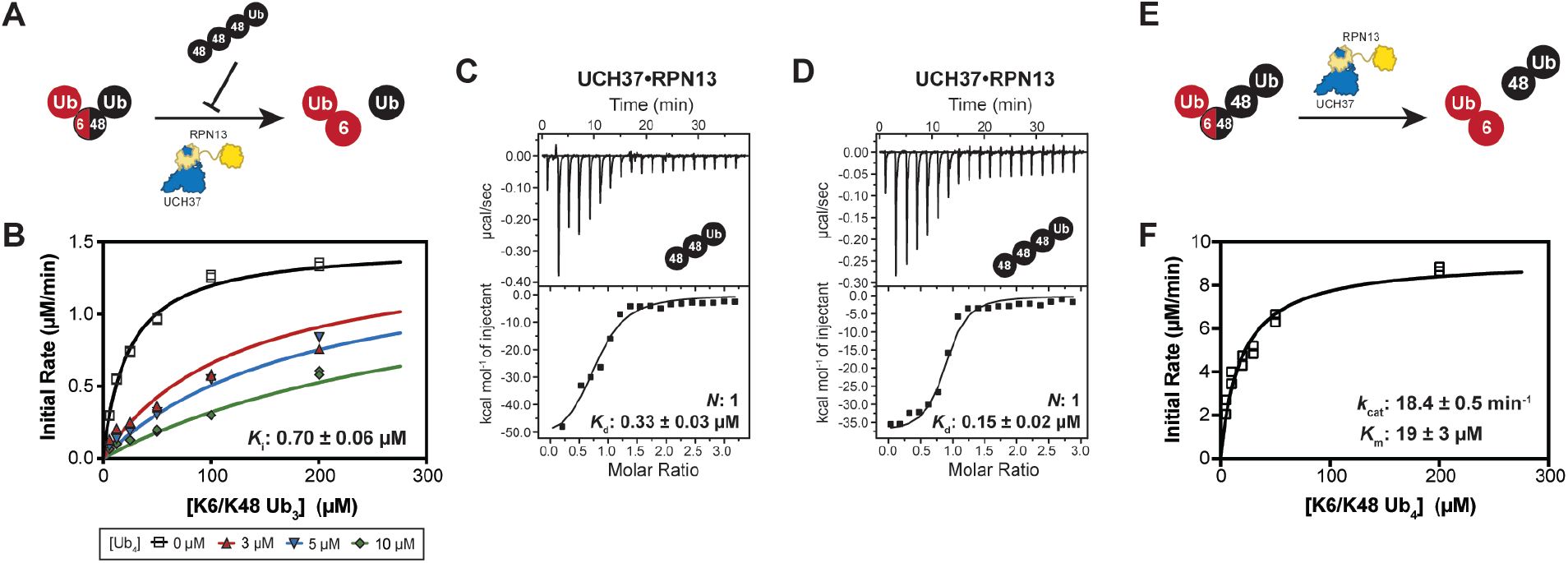
UCH37 Preferentially Binds K48 Chains. (A) Schematic for measuring the hydrolysis of native K6/K48 branched tri-Ub *in trans* with K48 tetra-Ub. (B) Michaelis-Menten plot for the hydrolysis of native K6/K48 branched tri-Ub by UCH37•RPN13 (0.5 μM) in the presence of K48 tetra-Ub. (C-D) ITC analysis of UCH37•RPN13 binding to K48-linked tri-Ub (C) and tetra-Ub (D). (E) Schematic for measuring the hydrolysis of native K6/K48 branched tetra-Ub *in cis*. (F) Michaelis-Menten plot for the hydrolysis of native K6/K48 branched tetra-Ub by UCH37•RPN13 (0.5 µM). All ITC curves are representative traces and reported *K*_d_s (C, D) are derived from averaging fits of two independent experiments. All kinetic curves (B, F) are averaged representative traces and constants are derived from averaging fits of independent experiments with SD (n = 2). See also Figure S4 and Table S1D.

The situation changes when additional K48 linkages are present *in cis*. A K6/K48 tetra-Ub chain was constructed by conjugating another Ub molecule to the distal K48 subunit of K6/K48 tri-Ub through a K48 linkage. This tetramer ostensibly has a homotypic K48 chain added *in cis* to a branched chain (Figure 4E). Kinetic analysis of the debranching reaction with the tetramer affords steady-state parameters that are similar to those obtained with K6/K48 tri-Ub (Figure 4F). The tetramer also binds with only two-fold higher affinity to UCH37•RPN13 (Figure S4F-G). These results suggest that unlike homotypic K48 chains presented *in trans*, *in cis* segments of K48-linked subunits do not inhibit the debranching reaction. Although future studies will be required to understand the molecular basis of UCH37•RPN13-catalyzed chain debranching, our kinetic analyses suggest that chains might be highly mobile once bound to UCH37 and discrimination between architectures could occur after the initial binding step.

### Proteasome-Bound UCH37 Debranches Chains

Since RPN13 recruits UCH37 to the proteasome, we wondered whether proteasome-bound UCH37 would also act as a debranching enzyme. Proteasomes (Ptsms) were purified from a HEK293 cell line stably expressing a HTBH-tagged (6xHis/TEV/biotin/6xHis) version of the proteasomal DUB RPN11 (HEK293^RPN11-HTBH^) (Figure 5A) (Wang et al., 2007). Western blot analysis shows the presence of UCH37 and RPN13 along with other 19S and 20S subunits (Figure 5B). We removed the other non-essential proteasomal DUB USP14 under high salt conditions (Besche et al., 2009; Lee et al., 2010) to ensure that the observed DUB activity can be ascribed to UCH37 (Figure S5A). Adding purified Ptsms deficient in USP14, but replete with UCH37 (herein referred to as wild-type Ptsm), to HMW K6/K48 chains results in a concentration-dependent formation of shorter chains along with mono-Ub (Figure 5C). As evidenced by intact MS, the formation of lower MW species is due to the removal of branch points (Figure 5D). A similar loss of branch points is observed with HMW K11/K48, and K48/K63 chains, indicating that Ptsm-catalyzed debranching is not idiosyncratic to K6/K48 chains (Figure S5C-E).

**Figure 5.**
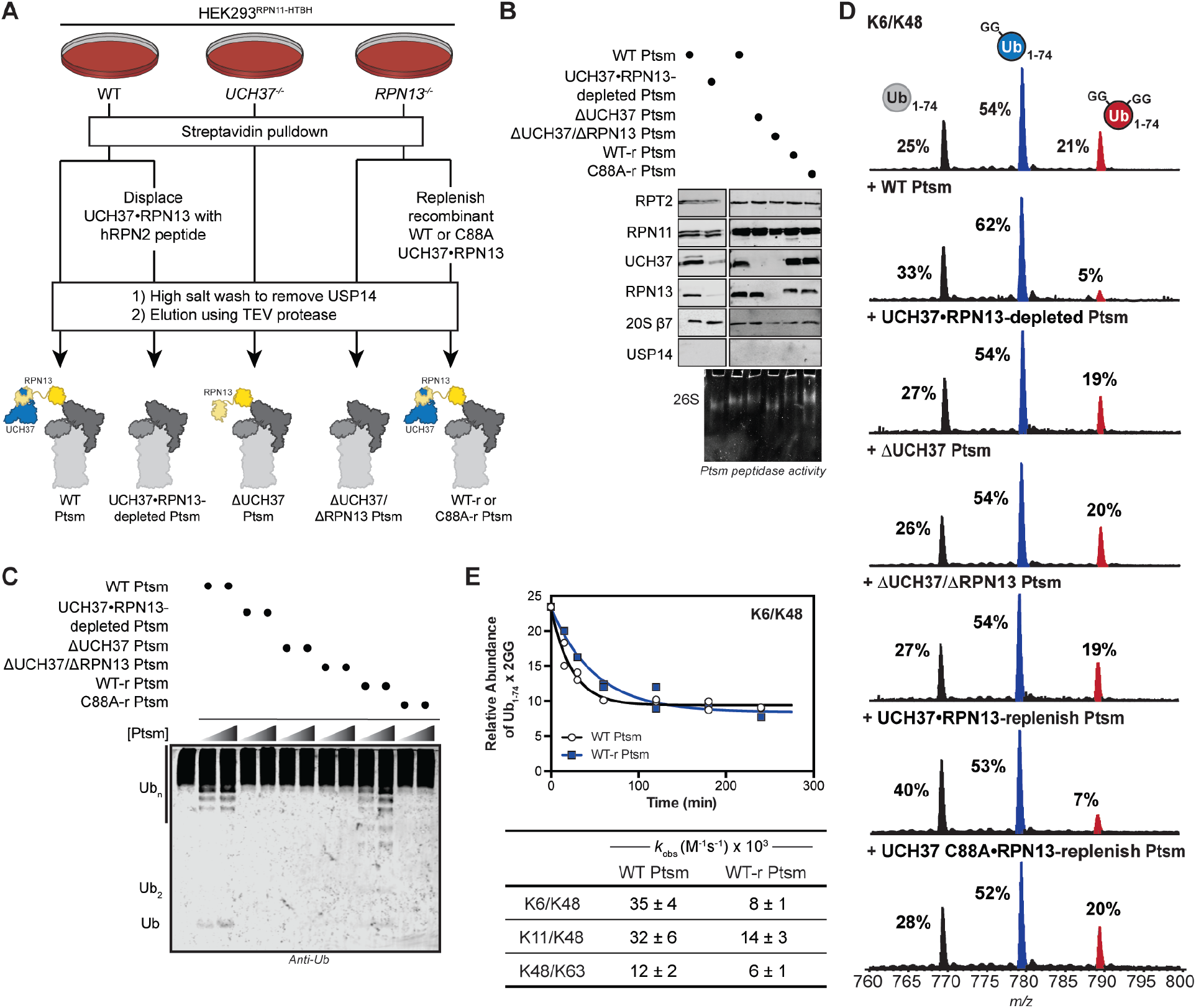
Proteasome-Bound UCH37 Debranches Ub Chains. (A) Purification scheme for the isolation of various human proteasome (Ptsm) complexes. (B) Western blot and Native PAGE (26S) analysis of purified Ptsm complexes (2 µg). (C) Western blot analysis of HMW K6/K48 chain debranching with increasing concentration of Ptsms (2 and 5 µg) using the α-Ub P4D1 antibody. (D) Ub MiD MS analysis of HMW K6/K48 chains subjected to each of the indicated Ptsm complexes (10 μg). Percentages correspond to the relative quantification values of the 11+ charge state for each Ub species: Ub_1-74_, 1xdiGly-Ub_1-74_, and 2xdiGly-Ub_1-74_. (E) Time course analysis of the removal of branch points from HMW K6/K48 chains by WT Ptsm (10 μg, black) and UCH37•RPN13-replenished Ptsm (10 μg, blue) (top). The 11+ charge state of the 2xdiGly-Ub_1-74_ species was used to measure the relative abundance of 2xdiGly-Ub_1-74_ at each time point. All curves are averaged representative traces from independent experiments with SD (n = 2). Table of kinetic parameters (*k*_obs_) obtained from fits to a first-order decay (bottom). Observed rate constants are normalized based on the concentration of UCH37. See also Figure S5 and Table S1E.

To then ascertain whether the debranching activity of purified Ptsms is due to UCH37, we used two orthogonal approaches. First, we displaced UCH37 and RPN13 from wild-type Ptsm complexes using a 38-mer peptide from the scaffolding subunit RPN2 (RPN2^916-953^) (Figure 5A) (Lu et al., 2015a). Subjecting the resulting UCH37•RPN13-depleted Ptsms to the fluorogenic Suc-LLVY-AMC peptide shows cleavage activity remains the same as wild-type proteasomes (Figure S5B). Ub-AMC hydrolysis, however, is severely compromised, supporting the notion that UCH37 is the only non-essential DUB on purified wild-type Ptsms (Figures S5B). Second, we used CRISPR/Cas9 genome editing to remove UCH37 from HEK293^RPN11-HTBH^ cells (Figure 5A). Ptsms purified from these cells are devoid of UCH37 (ΔUCH37 Ptsm), but still capable of cleaving Suc-LLVY-AMC (Figure S5B). Like UCH37•RPN13-depleted Ptsms, ΔUCH37 Ptsms display little activity toward HMW chains according to both western blot and intact MS (Figures 5C-D and S5C-E), suggesting UCH37 is responsible for the debranching activity.

For further corroboration, we initially tried to reconstitute ΔUCH37 Ptsms with active and inactive forms of UCH37. However, when active UCH37 was added back to ΔUCH37 Ptsms, the resulting complex displayed little activity toward Ub-AMC and HMW chains. As mentioned above, a similar problem was encountered when adding purified UCH37 to RPN13. Instead, we added back active and inactive recombinant UCH37•RPN13 complexes to RPN13-deficient Ptsms (Figure 5A). CRISPR/Cas9 was also used to delete RPN13 from HEK293^RPN11-HTBH^ cells. As shown by Native PAGE and western blot, RPN13-deficient Ptsms can be reconstituted with both active and inactive forms of UCH37•RPN13 (Figure 5B). Interrogating replenished Ptsms with HMW chains shows that chain debranching occurs with catalytically active UCH37, but not inactive UCH37 C88A (Figure 5C-D and S5C-E). Thus, we conclude that Ptsm-bound UCH37 debranches Ub chains.

### Kinetics of Proteasome-Bound UCH37 Resemble the UCH37•RPN13 Complex

Steady-state kinetic data with the UCH37•RPN13 complex showed that the rate of debranching occurs with a turnover number of 16-19 min^−1^, which means a K48 branch point can be removed within 3-4 s (1/*k*_cat_). Assuming the activity of the UCH37•RPN13 complex reflects the Ptsm-bound form, we would expect the kinetics of debranching to be commensurate with degradation rates, as several studies suggest it takes the proteasome tens of seconds to clear a ubiquitinated protein (Bard et al., 2019; Lu et al., 2015b; Peth et al., 2013). We tested this by determining the relative rates of debranching by UCH37•RPN13 and Ptsm-bound UCH37.

Middle-down MS was used to monitor the loss of branch points from HMW chains at different time points using UCH37•RPN13, wild-type Ptsms and active UCH37•RPN13-replenished Ptsms. These data show that branch points in K6/K48 and K48/K63 chains are rapidly depleted upon addition of UCH37•RPN13, wild-type Ptsm, and active UCH37•RPN13-replenished Ptsm (Figures S3E, 5E and S5H). After normalizing for differences in the concentration of UCH37, we find the observed rate constants are similar for UCH37•RPN13 and the Ptsm complexes (Figure S3E and 5E), suggesting the turnover number measured with the complex could reflect the activity of UCH37 on the proteasome.

While the overall reactivity patterns between UCH37•RPN13 and the proteasome complexes are largely the same with K6/K48 and K48/K63 chains, this is not true for K11/K48 chains. In the presence of UCH37•RPN13, the K11/K48 branch point is consumed within 15 min and there is little change in the unbranched portion of the chain even after 2 h (Figures S3E and S5G). With Ptsm complexes, the K11/K48 branch point is consumed rapidly, similar to reactions with UCH37•RPN13. After a slight delay, however, the unbranched portion also appears to decrease in abundance. Cleavage of both segments of the chain is dependent on the catalytic activity of UCH37. These results indicate that while UCH37•RPN13 and Ptsm-bound UCH37 display similar debranching kinetics, K11/K48 chains are dismantled to a greater extent in the presence of the Ptsm. As discussed in more detail below, this additional activity could be the result of proteasomal degradation.

### Chain Debranching Regulates Degradation

With data suggesting chain debranching occurs on a timescale relevant to degradation, we tested the impact of debranching on degradation. To do this, we generated a substrate modified with branched chains. While synthesizing K11/K48 chains, we found that the E2 fusion protein UBE2S-UBD (Bremm et al., 2010) modifies itself at three different positions: K117, K178, and K184 (Figure 6A). To determine whether branched chains are tethered to one of these sites, we appended a FLAG tag to UBE2S-UBD to separate the autoubiquitinated conjugates from unanchored chains. Middle-down MS analysis shows the extent of branching on UBE2S-UBD is nearly the same as in the bulk (∼9%), heterogenous mixture of HMW K11/K48 conjugates (Figure 6A). Thus, we decided to use polyubiquitinated UBE2S-UBD (K11/K48 Ub_n_-UBE2S-UBD) as a substrate for the Ptsm to measure the effect of debranching on degradation.

**Figure 6.**
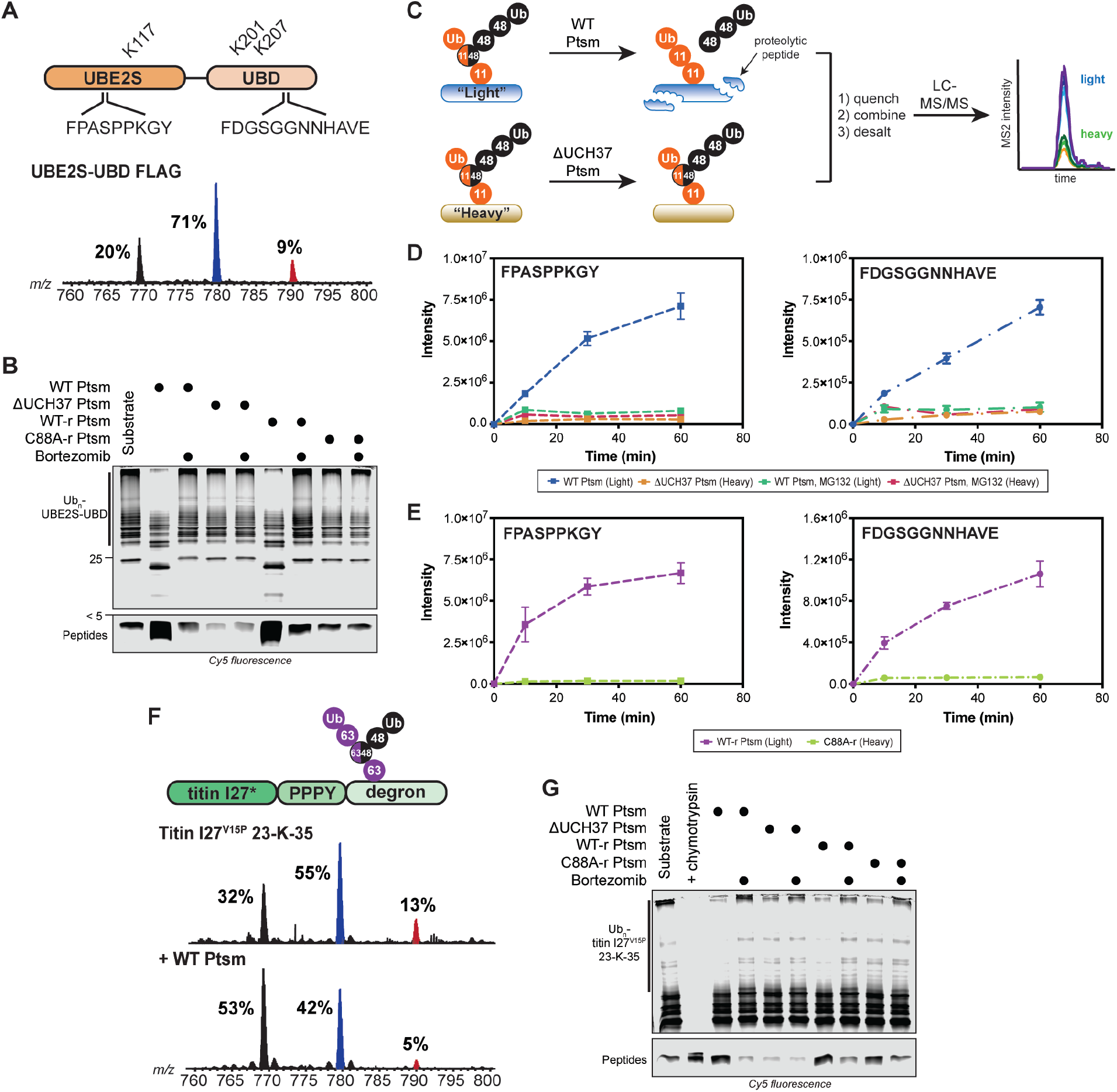
Ubiquitin Chain Debranching Promotes Proteasomal Degradation. (A) UBE2S-UBD construct (top) showing ubiquitination sites (K117, K201, and K207) and the proteasome-derived UBE2S-UBD peptides detected by parallel reaction monitoring (PRM) MS. Ub MiD MS analysis of K11/K48-Ub_n_-UBE2S-UBD-FLAG (bottom). (B) Degradation of K11/K48-Ub_n_-UBE2S-UBD using the indicated Ptsm complexes (5 µg) under multi-turnover conditions in the presence and absence of bortezomib (1 μM). (C) Schematic for tracking the formation of specific transition ions of UBE2S-UBD-derived proteolytic peptides. All reactions are completed as a two-plex experiment using ubiquitinated light and heavy (^15^N) UBE2S-UBD with different Ptsm complexes. (D-E) PRM MS analysis of UBE2S-UBD peptides formed by Ptsms. The UBE2S peptide FPASPPKGY is shown on the left and the UBD peptide FDGSGGNNHAVE is on the right. (D) Light and heavy UBE2S-UBD were mixed with either WT Ptsm (5 μg) or ΔUCH37 Ptsm (5 μg), respectively, in the presence and absence of MG132 (10 μM). (E) Light and heavy UBE2S-UBD were mixed with either WT UCH37•RPN13-replenished Ptsm (WT-r Ptsm; 5 μg) or UCH37 C88A•RPN13-replenish Ptsm (C88A-r Ptsm; 5 μg). (F) K48/K63-Ub_n_-titin-I27^V15P^-23-K-35 construct (top) used in this study. Ub MiD MS analysis of K48/K63-Ub_n_-titin-I27^V15P^-23-K-35 (middle) followed by treatment with WT Ptsm (10 μg, bottom). (G) Degradation of K48/K63-Ub_n_-titin-I27^V15P^-23-K-35 using the indicated Ptsm complexes (5 µg) under multi-turnover conditions in the presence and absence of bortezomib (1 μM). Ub MiD MS percentages correspond to the relative quantification values of the 11+ charge state for each Ub species: Ub_1-74_, 1xdiGly-Ub_1-74_, and 2xdiGly-Ub_1-74_. Multiple-turnover assays were tracked by tricine-SDS-PAGE and visualized by cy5 fluorescence. All MS curves are representative depictions from the sum of the transition ions of each monitored peptide (D-E) with a dashed line connecting the averaging of independent experiments with SD (n = 4). See also Figure S6 and Table S1F-G.

K11/K48 Ub_n_-UBE2S-UBD was fluorescently labeled using a cysteine-reactive Cy5 dye and degradation by different Ptsm complexes was monitored by SDS-PAGE under multi-turnover conditions. The HMW bands corresponding to K11/K48 Ub_n_-UBE2S-UBD disappear in the presence of both wild-type Ptsm and UCH37•RPN13-replenished Ptsm, but not with ΔUCH37 or UCH37 C88A•RPN13-replenished Ptsms (Figure 6B). The appearance of peptides also depends on the activity of UCH37.

To confirm that UBE2S-UBD-derived peptides are indeed produced, we monitored Ptsm-mediated peptide formation using parallel reaction monitoring (PRM) MS (Figure 6C). Two peptides from separate regions of UBE2S-UBD–one from UBE2S (FPASPPKGY) and another from the UBD (FDGSGGNNHAVE)–could be detected reproducibly. Both peptides are products of the Ptsm, as inhibition with MG132 abolishes their formation. We then directly compared the abundance of peptides at various time points with different Ptsm complexes by treating the ‘light’ form of K11/K48 Ub_n_-UBE2S-UBD with one Ptsm complex and the ‘heavy’ form (K11/K48 Ub_n_-^15^N-UBE2S-UBD) with another and mixing the resulting peptides. Labeling UBE2S-UBD with ^15^N does not affect the nature of chains it attaches to itself (Figure S6B). The PRM results show that peptide formation is more robust with wild-type Ptsms compared to reactions with ΔUCH37 Ptsms (Figures 6D and S6C). This difference can be ascribed to the activity of UCH37 since peptides accumulate to a greater extent when Ptsms contain active UCH37, but not inactive UCH37 C88A (Figure 6E and S6D). Moreover, if branch points are removed from K11/K48 Ub_n_-UBE2S-UBD prior to the addition of Ptsms, ΔUCH37 Ptsms are now capable of degrading UBE2S-UBD (Figure S6F), demonstrating that these Ptsms are indeed active. Together, these results suggest chain debranching by UCH37 is important for promoting degradation.

To assess whether the degradation of other branched chain-modified substrates depends on the activity of UCH37, we availed a well-established model substrate for Ptsm degradation—titin-I27^V15P^-23-K-35 (Bard et al., 2019). Titin-I27^V15P^-23-K-35 contains a destabilizing V15P substitution and a C-terminal initiation region with a single lysine for Ub chain attachment. We envisioned building K48/K63 branched chains on the C-terminal lysine by first installing K63 chains using the yeast E3 Rsp5 and then adding K48 branch points using UBE2R1. As shown by middle-down MS, this approach works well and wild-type Ptsms are able to debranch the K48/K63 chains attached to titin-I27^V15P^-23-K-35 (Figure S6F). The MS data also reveals a slight reduction in the unbranched portion similar to what we observed with K11/K48 chains, suggesting cleavage of the unbranched portion correlates with proteasomal degradation (Figure 6F).

Degradation of K48/K63 Ub_n_-titin-I27^V15P^-23-K-35 was monitored by SDS-PAGE and visualized by labeling cysteine-135 of titin-I27^V15P^-23-K-35 with Cy5. As with K11/K48 Ub_n_-UBE2S-UBD, wild-type Ptsms deplete the K48/K63 Ub_n_-titin-I27^V15P^-23-K-35 conjugates and catalyze robust peptide formation (Figure 6G). ΔUCH37 Ptsms, by contrast, do not reduce the abundance of polyubiquitinated species or generate peptide products. A comparison of the reactions with replenished Ptsms also shows that the polyubiquitinated species are consumed and peptides are produced to a greater extent with active UCH37. Thus, the ability of UCH37 to promote degradation by removing branch points applies to different substrates and different K48 branched chains.

### UCH37 Potentiates Proteasomal Degradation In Cells

Based on our *in vitro* data, we would expect the loss of UCH37 to impede degradation in cells. To test this hypothesis, we initially measured differences in the stability of the GFP^u^ proteasome activity reporter (Bence et al., 2001) in the presence and absence of UCH37 (Figure 7A). A recent study showed that GFP^u^ is a substrate of the endoplasmic reticulum-associated degradation (ERAD) pathway and modified with K29/K48 branched chains (Leto et al., 2019). Flow cytometry shows that the levels of GFP^u^ nearly double in UCH37-deficient HEK293 cells relative to wild-type. Although GFP^u^ levels do not increase to the same degree as complete proteasome inhibition (Figure 7B), these data support the supposition that UCH37 potentiates degradation.

**Figure 7.**
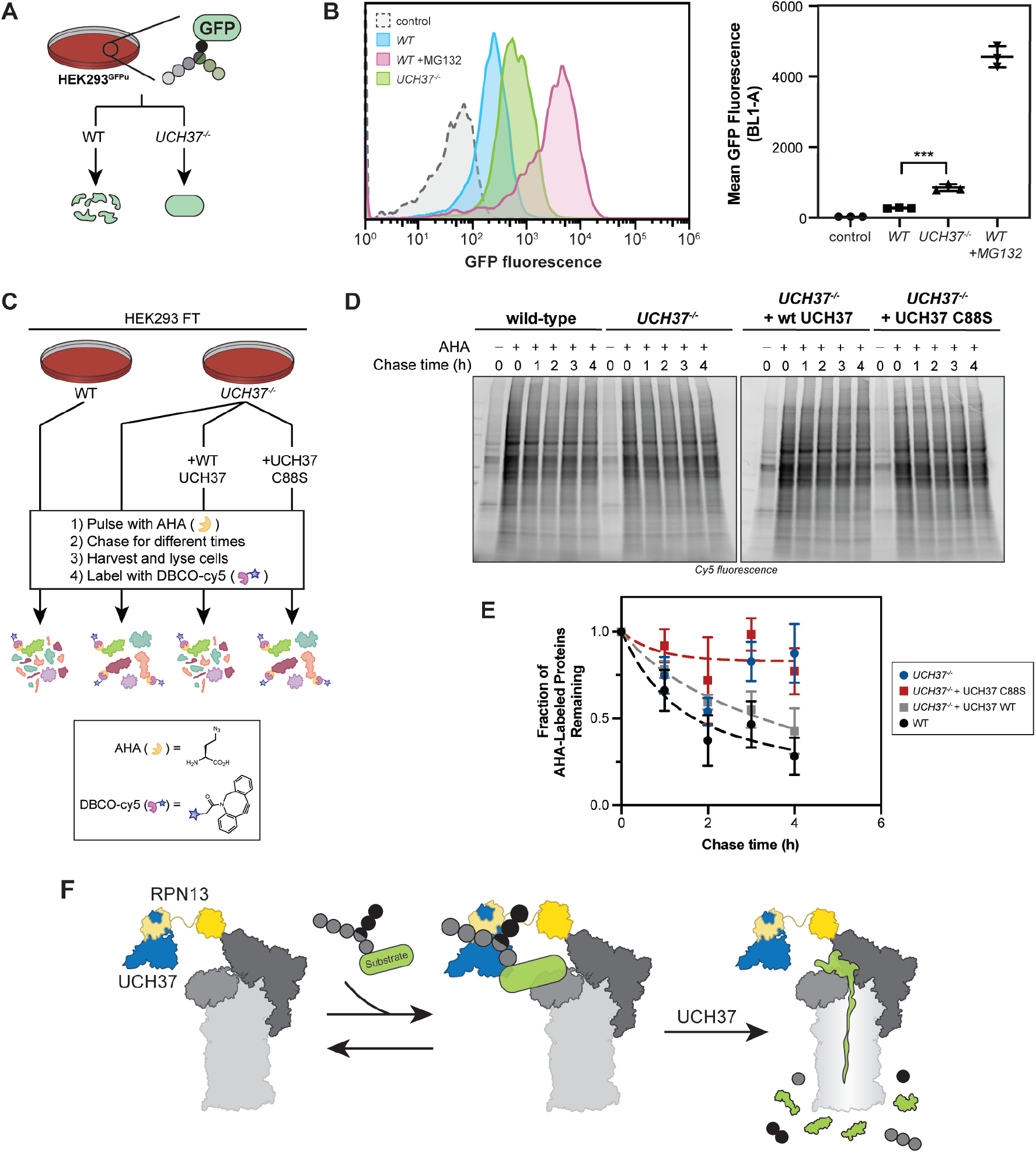
UCH37 Potentiates Proteasomal Degradation in Cells. (A) Schematic for GFP^u^ turnover in cells. (B) Fluorescence histograms of control, wild-type (treated with and without MG132), and *UCH37^−/−^* GFP^u^ cells measured using flow cytometry. Quantitation of mean GFP fluorescence from averaging fits of three independent experiments. \*\*\**P*<0.0005 (Student’s t-test). (C) Schematic for measuring global protein turnover using azidohomoalanine (AHA) labeling of newly synthesized proteins in cells. (D) Cy5 fluorescence SDS-PAGE analysis of the turnover of AHA-labeled proteins at different chase times in wild-type, *UCH37^−/−^*, and *UCH37^−/−^* HEK293 FT cells expressing WT UCH37 or catalytically inactive UCH37 C88S. (E) Densitometric quantification of AHA labeled proteins around 50 kDa relative to *t* = 0. Results are shown as mean ± s.e.m from three individual experiments. (F) Model showing UCH37-catalyzed Ub chain debranching promotes proteasomal degradation. AHA incorporated proteins were labeled with DBCO-Cy5, separated by SDS-PAGE and visualized by Cy5 fluorescence. See also Figure S7.

Next, we wanted to assess the impact of UCH37 on global protein turnover. Pulse-chase experiments were performed on a proteome-wide scale. Newly synthesized proteins were labeled with the unnatural amino acid azidohomoalanine (AHA) (Figure 7C) (McShane et al., 2016). Cells were then chased in AHA-free medium for different lengths of time. At each time point, AHA-labeled proteins were visualized after conjugation to a Cy5 dye using strain-promoted click chemistry. As shown with wild-type HEK293 cells, AHA-labeled proteins decrease in abundance over time, and proteasome inhibition with MG132 prevents their loss (Figure S7A). In cells lacking UCH37, the amount of AHA-labeled proteins remains relatively constant (Figure 7D). Degradation can then be rescued only when null cells are complemented with active UCH37 (Figure 7D-E). Together, the biochemical and cellular data have led us to conclude that UCH37 acts as an important regulator of proteasomal degradation by debranching Ub chains (Figure 7F).

## DISCUSSION

In this study, we identify UCH37 as a DUB capable of debranching Ub chains. The remarkable topological specificity of UCH37 occurs in conjunction with K48 linkage selectivity. When UCH37 is bound to RPN13 alone or in context of the proteasome, the rate of debranching is dramatically enhanced. This activity has surprising consequences on protein turnover, as our work reveals that UCH37 removes branch points to enhance proteasomal degradation.

The prevailing view is that UCH37 contributes to the substrate selection process by trimming Ub chains to reduce the residence time on the proteasome (Collins and Goldberg, 2017; de Poot et al., 2017). This supposition has largely been based on the premise that UCH37 cleaves homotypic K48 chains. The kinetics of homotypic K48 chain cleavage by UCH37, however, are exceedingly slow even in the presence of RPN13 (Bett et al., 2015; Lu et al., 2017; Yao et al., 2006). If the purpose is to rescue poorly ubiquitinated proteins from the fate of the proteasome, then UCH37 would have to act quickly. Take for example, the other non-essential cysteine-dependent DUB that associates with the proteasome—USP14. Evidence suggests that USP14 functions before the commitment step (Lee et al., 2016), which does not occur until an unstructured region of the substrate is inserted into the AAA+ ATPase motor of the 19S regulatory particle and the proteasome adopts a conformation conducive to proteolysis (Bard et al., 2019; Lu et al., 2015b). These steps are estimated to take only a few seconds, which means USP14 operates on a much faster timescale and UCH37 would have to as well in order to oppose the degradation process.

By identifying K48 branched chains as targets, we can develop a better picture of how UCH37 fits into the kinetic scheme of degradation. Steady-state kinetic analyses suggest that UCH37 in complex with RPN13 selectively removes K48 branch points every ∼3 s (1/*k*_cat_) and the level of activity on the proteasome is similar. Thus, in light of the detailed kinetics that have been reported for degradation (Bard et al., 2019; Lu et al., 2015b), Ub chain debranching would occur around the same time as either the commitment step or deubiquitination by the intrinsic metalloprotease RPN11/POH1. Based on this logic, UCH37 would not have a chance to debranch chains prior to the commitment step.

How does Ub chain debranching stimulate degradation? The lid of the proteasome houses multiple Ub receptors (RPN13, RPN10, and RPN1) providing a versatile recognition platform for substrates modified with different Ub chains (Deveraux et al., 1994; Husnjak et al., 2008; Schreiner et al., 2008; Shi et al., 2016). Given the multivalency and high local concentration of Ub in branched chains, cooperation between receptors could promote high avidity interactions. According to a recent study, it may not be necessary for multiple receptors to engage branched chains since RPN1 binds K11/K48 branched chains with higher affinity than the linear counterparts (Boughton et al., 2020). Branched chains could thus prioritize substrates for the proteasome at the initial binding stage. However, affinity does not always correlate with degradation efficiency. Longer chains, for example, bind more tightly to the proteasome than shorter chains, yet degradation rates for substrates modified with either long or short chains are nearly the same (Martinez-Fonts et al., 2020).

Debranching could be necessary to take full advantage of the avidity afforded by branched chains yet still ensure efficient degradation. It is possible that branched chains bind in a configuration that impedes the proteasome from transitioning into a catalytically competent conformation or RPN11-mediated deubiquitination (Hyoung et al., 2007; Kim et al., 2009). The removal of branch points could mitigate this problem. Another possibility is that branched chains could slow down product release. Debranching would then free the proteasome for additional rounds of proteolysis.

The ability of a DUB to selectively target branch points for removal is unprecedented. How UCH37 selects K48 branch points from a sea of unbranched chains is unclear. Considering UCH37 engages both branched and unbranched K48 chains, there must be a mechanism for UCH37 to transfer between different segments of a chain. Understanding the molecular details of debranching could provide insight into the design of effective inhibitors of UCH37, the development of which is even more pressing given its role in promoting proteasomal degradation—a process many cancer cells rely on for survival (Deshaies, 2014).

## AUTHOR CONTRIBUTIONS

K.K.D., S.O.C., J.D., H.B. and R.G.G. conducted the experiments. E.R.S. supervised the experiments. K.K.D., S.O.C., J.D., H.B, and E.R.S. designed the experiments and wrote the manuscript.

## ACKNOWLEDGEMENTS

We thank Dr. Steve Eyles (UMass Amherst) for assistance with high-resolution mass spectrometry. This work was funded by the NIH (RO1GM110543 to E.R.S.), a NSF Graduate Research Fellowship (GRFP1451512 to K.K.D.), and a NIH Chemistry and Biology Training Grant (T32GM008515 to J.D. and H.B.). The data described herein were acquired on an Orbitrap Fusion mass spectrometer funded by National Institutes of Health grant 1S10OD010645-01A1.

## MATERIALS

### Antibodies

Anti UCH37 (Abcam, Cat. # ab124931)

Anti RPN11 (Abcam, Cat. # ab109130)

Anti RPN13 (Cell Signaling Technology, Cat. # D9Z1U)

Anti RPT2 (Abcam, Cat. # ab3317)

Anti PSMB7 (R&D Systems, Cat. #MAB7590)

Anti USP14 (Abcam, Cat. # ab56210)

Anti Ub (P4D1, Enzo Lifesciences, Cat. # BML-PW0930)

Anti K48-linkage Specific (Cell Signaling Technology, Cat. # D9D5)

Goat Anti Mouse IR Dye 800CW (LI-COR Biosciences, Cat. # 926-32210)

Goat Anti Rabbit IR Dye 680RD (LI-COR Biosciences, Cat. # 926-68071)

Goat Anti Rabbit IR Dye 800CW (LI-COR Biosciences, Cat. # 926-32211)

### Bacterial and Viral Strains

Rosetta 2(DE3)pLysS (EMD Millipore Novagen, Cat. # 71403-3)

BL21(DE3)pLysS (Promega, Cat. # L1191)

One Shot™ TOP10 (Fisher Scientific, Cat. # C404003)

One Shot™ Stbl3 (Fisher Scientific, Cat. # C737303)

### Chemicals, Peptides, and Recombinant Proteins

ProBlock Gold Mammalian Protease Inhibitor Cocktail (Gold Biotechnology, Cat. # GB-331-5)

Simple Stop™ 2 Phosphatase Inhibitor Cocktail (Gold Biotechnology, Cat. # GB-451)

Ammonium Chloride (^15^N, 99%) (Cambridge Isotope Laboratories, Cat. # NLM-467)

SYPRO Ruby Stain (Fisher Scientific, Cat. # S12000)

Cy5 maleimide (Lumiprobe, Cat. #23080)

DBCO-Cy5 (Sigma, Cat. # 777374)

Click-iT™ AHA (L-Azidohomoalanine) (Fisher Scientific, Cat. #C10102)

Trypsin (Promega, Cat. # V5113)

Chymotrypsin (Promega, Cat. # V1061)

Formic Acid (Sigma-Aldrich, Cat. # 399388)

Acetic Acid (Fisher Scientific, Cat. # 351269-4)

Creatine phosphate disodium salt (Abcam, Cat. # ab146255)

Creatine Kinase (Sigma-Aldrich, Cat. # 10127566001)

Adenosine-5’-triphosphate (Goldbio, Cat. # A-081-5)

Ub-AMC (Boston Biochem, Cat. # U-550)

Suc-LLVY-AMC (Boston Biochem, Cat. # S-280)

MG132 (Fisher Scientific, Cat. # 508339)

Bortezimib (Selleck Chemicals, Cat. # S1013)

Polybrene (Sigma, Cat. # TR-1003-G)

Lipofectamine 3000™ (Fisher Scientific, Cat. # L3000008)

Doxycycline hyclate (Sigma, Cat. # D9891)

AQUA peptides (Cell Signaling Technology, see Table S1)

### Critical Commercial Assays

Alt-R^®^ S.p. Cas9 Nuclease 3NLS (Integrated DNA Technologies, Cat. # 1081058)

PTMScan^®^ Ubiquitin Remnant Motif (Cell Signaling Technology, Cat. # 14482)

pENTR™/SD/D-TOPO™ Cloning Kit (Fisher Scientific, Cat. # K242020)

Gateway LR Clonase II Enzyme Mix (Fisher Scientific, Cat. # 11-791-020)

### Experimental Models: Cell Lines

HEK293 Expressing Rpn11-HTBH (Applied Biological Materials, T6007)

HEK293 FT (ATCC, CRL-3216)

HEK293 GFPu-1 (ATCC, CRL-2794)

### Oligonucleotides

CRISPR KO sgDNA sequence UCH37 (1): GTTACTGAACTGTACCCACC

CRISPR KO sgDNA sequence UCH37 (2): CGCCTAAATGGACATCCTGG

CRISPR KO sgDNA sequence RPN13: CACGAACTCTCTGCGCTAGG

### Recombinant DNA (Source, Identifier)

pMCSG20: NleL (aa 170-782) (Valkevich et al., 2014, N/A)

pQE30: SortaseΔN25 (SrtA) (Crowe et al., 2016, gifted from O. Schneewind)

pVP16: UCH37 (DNASU, HsCD00084019)

pET19: RPN13 or Adrm1 (Addgene, Plasmid #19423)

pGEX-6P1: hRPN2 (aa 916-953) (Lu et al., 2015; gifted from K. Walters)

pET28b: E1 (Trang et al., 2012; N/A)

pGEX-4T2: UBE2D3 (Valkevich et al., 2014, N/A)

pGEX-6P1: UBE2S-UBD (Addgene, Plasmid #66713)

pGEX-6P1: AMSH (aa 219-424) (Trang et al., 2012, N/A)

pOPINK: OTUD1 (Addgene, Plasmid #61405)

pVP16: OTUB1 (Pham et al., 2015, N/A)

pOPINS: UBE3C (Addgene, Plasmid #66711)

pDEST17: UBE2R1 (Pham et al., 2015, N/A)

pST39: UBE2N/UBE2V2 (Pham et al., 2015, N/A)

pET28b: UBC1 (DNASU, ScCD00009212)

pOPINK: RSP5 (DNASU, ScCD00008707)

pOPINS: Titin I27^V15P^ 23-K-35 (Bard et al., 2019, N/A)

pET22b: Ub and Ub variants (Valkevich et al., 2012, N/A)

pET28b: SUMO2 (Addgene, Plasmid #25102)

pET22b: GFP (Addgene, Plasmid #11938

pMD2.G (Addgene, Plasmid #12259)

psPAX2 (Addgene, Plasmid #12260)

pINDUCER21 (Addgene, Plasmid #46948)

### Software and Algorithms

Typhoon FLA 9500 (GE Healthcare)

Odyssey CLx Imager (LICOR)

Image Studio software (LICOR Biosciences

Prism 8 (Graphpad Software)

OriginLab 7 SR4 (OriginLab Corporation)

Xcalibur 3.0 (Thermo Fisher Scientific)

Pinpoint 1.4 (Thermo Fisher Scientific)

Proteome Discoverer 2.3 (Thermo Fisher Scientific)

Mash Suite (Guner et al., 2014)

FlowJo 10.4 (FlowJo, LLC)

### Other

His60 Ni Superflow resin (Clontech, Cat. # 635660)

Glutathione resin (GenScript, Cat. # L00206)

Amylose resin (NEB, Cat. # E8021S)

Streptavidin resin (GenScript, Cat. # L00353)

Anti-Flag M2 Affinity Gel (Sigma, Cat. # A2220)

Slide-A-lyzer MINI dialysis units (3.5 kDa MWCO) (Thermo Scientific, Cat. # PI69552)

100mg SEP-PAK C18 column (Waters, Cat. # wat043395)

C18 StageTips (Thermo Scientific, Cat. # SP301)

Zeba Spin Desalting Column (Thermo Scientific, Cat. # 89889)

NuPAGE Novex 12% Bis-Tris Protein Gels (Fisher Scientific, Cat. # NPO343BOX)

4-20% Mini-PROTEAN Gels (Bio-Rad, Cat. # 4561096)

Syringe Filters, PES, 0.45 µm (Genesee Scientific, Cat. # 25-246)

## CONTACT FOR REAGENT AND RESOURCE SHARING

Further information and requests for reagents may be directed to and will be fulfilled by Eric Strieter

## EXPERIMENTAL MODEL AND SUBJECT DETAILS

### Human Cells

HEK293 cells stably expressing His-biotin affinity tagged human RPN11 (RPN11-HTBH) (Wang et al., 2007), HEK293FT, and HEK293 GFPu-1 cells were cultured at 37°C under 5 % CO2 using high glucose DMEM supplemented with 10% FBS, 1xGlutamax (Gibco), and 1xPen/Strep.

## METHOD DETAILS

### Generation of CRISPR KO Cell Lines

Guide RNAs were designed for UCH37 and RPN13 using design tools from Harvard and the Broad Institute. The gRNAs were purchased from IDT to be used in their Alr-R® CRISPR-Cas9 system. CRISPR reactions were performed according to the protocol provided by IDT. Briefly, a 1:1 annealed complex of gRNA: tracrRNA was prepared by mixing 1 μM of each RNA in nuclease-free duplex buffer and warming to 95°C for 5 min before cooling to room temperature. Once the annealed complex was prepared, a ribonucleotide-protein complex (RNP) was prepared by mixing the RNA complex with Alt-R® S.p. Cas9 Nuclease in Opti-MEM media and incubating at room temp for 5 min. A transfection containing the RNP mixture was then prepared by diluting the RNP in Opti-MEM, adding Lipofectamine RNAimax, briefly vortexing, and allowing the mixture to incubate for 20 min at room temperature. The transfection mixture was then added to a 48 well plate followed by 80,000 cells in antibiotic free media. These cells were allowed to grow for 2 days before trypsinization and dilution to single cell per well density in 96 well plates. Single colonies were identified and screened for protein expression by western blot using appropriate antibodies. Colonies that showed loss of UCH37 or RPN13 were further screened by sequencing and analysis using TIDE software. Colonies with confirmed Indels were used in future experiments.

### Lentivirus Packaging, Infection and Cell Line Creation

HEK293 FT packaging cells were seeded in 60 mm dishes four days prior to transfection at a density of 200,000 cells. HEK293 FT cells were transiently transfected with a mix of 3 µg packaging vector (psPAX2), 2 µg envelope vector (pMD2.G), and 3 µg pINDUCER21 vector containing the ORF of either WT UCH37 or C88S UCH37 in Opti-MEM (Gibco) using Lipofectamine 3000 following the recommended protocol. At 48 h post-transfection, the cell culture media was collected and filtered through a 0.45 µm filter. 9 mL of lentivirus-containing supernatant was combined with 9 mL of DMEM containing 7.2 µg polybrene. *UCH37^−/−^* HEK293 FT cells were seeded in T25 flasks at a density of 420,000 cells and incubated under standard cell culture conditions until 60% confluency was reached. A total of 18 mL of lentivirus was used for infection where 6 mL of lentivirus was introduced to cells every 12 h for a total of 36 h infection. After the last infection, the supernatant was removed, and cells were grown in fresh DMEM for 48 h. Infected cells were selected by flow cytometry using the BD FACSAria II SORP cell sorter (BD Biosciences) equipped with 355 (UV), 405, 488, 561, and 640 nm lasers with 16 color analysis capabilities for the detection of high eGFP expression in the target cells.

### Protein Purification

All plasmids used for purification are described in the key resources table. Proteins were purified at 4°C unless otherwise indicated. After the final chromatography step, all proteins were concentrated in Amicon Ultra spin concentrators, aliquoted, flash frozen in liquid N2 and stored at −80 °C. Concentrations of all proteins were determined by BCA assay.

### Purification of Ubiquitination machinery

E1, UBE2D3, UBE2R1, UBE2N/UBE2V2, UBC1, and UBE3C were purified as previously described (Trang et al., 2012; Pham et al., 2015; Bashore et al., 2015; Michel et al., 2015). Briefly, E1, UBE2D3, UBE2R1, and UBE2N/UBE2V2 constructs were expressed in Rosetta 2(DE3)pLysS *E.coli* cells in LB media supplemented with appropriate antibiotics at 37°C to OD600 ∼0.6-0.8 and transferred to 18°C for 16 h after induction with IPTG. Cultures were harvested, resuspended in lysis buffer A (50 mM Tris pH7.5, 300 mM NaCl, 1 mM EDTA and 10 mM imidazole), lysed by sonication, and clarified by centrifugation. Clarified lysate was then incubated with Ni-NTA resin for 1 h, washed with lysis buffer A, and eluted into Ni-NTA elution buffer (lysis buffer A plus 300 mM imidazole).

NleL (aa 170-782) were purified as previously (Valkevich et al., 2014). Briefly, NleL was expressed in BL21(DE3)pLysS *E.coli* cells in LB media supplemented with appropriate antibiotics at 37°C to OD600 ∼0.6 and transferred to 16°C for 16 h after induction with IPTG. Cultures were harvested, resuspended in lysis buffer B (50 mM Tris pH8.0, 200 mM NaCl, 1 mM EDTA and 1 mM DTT), lysed by sonication, and clarified by centrifugation. Clarified lysate was then incubated with GST resin for 1 h, washed with lysis buffer B, and eluted into GST elution buffer (lysis buffer B plus 10 mM reduced glutathione). Eluate was concentrated in TEV protease buffer (50 mM Tris pH8.0, 150 mM NaCl, and 0.5 mM TCEP), cleaved overnight with TEV protease, and further purified using anion exchange chromatography.

UBE2S-UBD was purified as previously described (Bremm et al., 2010). Briefly, UBE2S-UBD constructs were expressed in Rosetta 2(DE3)pLysS *E.coli* cells in LB media supplemented with appropriate antibiotics at 37°C to OD600 ∼0.6-0.8 and transferred to 16°C for 16 h after induction with IPTG. Cultures were harvested, resuspended in lysis buffer C (270 mM sucrose, 50 mM Tris pH8.0, 50 mM NaF, and 1 mM DTT), lysed by sonication, and clarified by centrifugation. Clarified lysate was then incubated with GST resin for 1 h, washed with high salt buffer A (25 mM Tris pH8.0, 500 mM NaCl, and 5 mM DTT), followed by low salt buffer A (25 mM Tris pH8.0, 150 mM NaCl, and 5 mM DTT), and resuspended in 3C protease buffer (50 mM Tris pH8.0 and 150 mM NaCl) for on-resin cleavage with HRV 3C protease overnight.

Minor modifications were made for the expression of ^15^N-UBE2S-UBD. Using the high-cell-density expression method adapted from Marley *et al*. (2001), cultures were grown to an OD600 of 0.8 at 37°C, concentrated (4x), and transferred into M9 media. Cells were incubated for 1 h to allow for discharge of unlabeled metabolites and then supplemented with 20% w/v glucose and 0.5 g ^15^NH4Cl, induced with IPTG, and grown overnight as described above.

RSP5 were purified as previously described (Worden et al., 2017). Briefly, RSP5 was expressed in Rosetta 2(DE3)pLysS *E.coli* cells in 2xYT media supplemented with appropriate antibiotics at 37°C to OD600 ∼0.8 and transferred to 18°C for 16 h after induction with IPTG. Cultures were harvested, resuspended in lysis buffer D (50 mM HEPES pH7.5, 250 mM NaCl, 1 mM EDTA, 1 mg/mL lysozyme, 1X protease cocktail and 5 mM PMSF), lysed by sonication, and clarified by centrifugation. Clarified lysate was then incubated with GST resin for 1 h, washed with high salt buffer B (25 mM HEPES pH8.0 and 500 mM NaCl), followed by low salt buffer B (25 mM HEPES pH8.0 and 150 mM NaCl), and resuspended in 3C protease buffer for on-resin cleavage with HRV 3C protease overnight. Eluate was concentrated and run on a Superdex 200 (GE) gel filtration column in 50 mM HEPES pH7.5, 50 mM NaCl, and 5% glycerol.

### Purification of Deubiquitinating Enzymes

OTUB1 was purified as previously described (Pham et al., 2015). Briefly, OTUB1 was expressed in Rosetta 2(DE3)pLysS *E.coli* cells in LB media supplemented with appropriate antibiotics at 37°C to OD600 ∼0.6-0.8 and transferred to 18°C for 16 h after induction with IPTG. Cultures were harvested, resuspended in lysis buffer A, lysed by sonication, and clarified by centrifugation. Clarified lysate was then incubated with Ni-NTA resin for 1 h, washed with lysis buffer A, and eluted into Ni-NTA elution buffer.

AMSH was purified as previously described (Trang et al., 2012). Briefly, AMSH was expressed in BL21(DE3)pLysS *E.coli* cells in LB media supplemented with appropriate antibiotics at 37°C to OD600 ∼0.6 and transferred to 16°C for 16 h after induction with IPTG and addition of ZnCl2. Cultures were harvested, resuspended in lysis buffer E (50 mM Tris pH7.5, 300 mM NaCl and 2 mM DTT), lysed by sonication, and clarified by centrifugation. Clarified lysate was then incubated with GST resin for 1 h, washed with high salt buffer A, followed by low salt buffer A, and resuspended in 3C protease buffer for on-resin cleavage with HRV 3C protease overnight.

OTUD1 was purified as previously described (Mevissen et al., 2013). OTUD1 was expressed in Rosetta 2(DE3)pLysS *E.coli* cells in LB media supplemented with appropriate antibiotics at 37°C to OD600 ∼0.6-0.8 and transferred to 20°C for 16 h after induction with IPTG. Cultures were harvested, resuspended in lysis buffer C, lysed by sonication, and clarified by centrifugation. Clarified lysate was then incubated with GST resin for 1 h, washed with high salt buffer A, followed by low salt buffer A, and resuspended in 3C protease buffer for on-resin cleavage with HRV 3C protease overnight. Eluate was concentrated and purified using anion exchange chromatography.

### Purification of hRPN2 peptide

hRPN2 (aa 916-953) construct was expressed in BL21(DE3)pLysS *E.coli* cells in LB media supplemented with appropriate antibiotics at 37°C to OD600 ∼0.5 and transferred to 18°C for 16 h after induction with IPTG. Cultures were harvested, resuspended in lysis buffer E containing 1 mM PMSF, lysed by French press, and clarified by centrifugation. Clarified lysate was then incubated with GST resin for 2 h, washed with high salt buffer A, followed by low salt buffer A, and eluted into GST elution buffer (lysis buffer E plus 10 mM reduced glutathione). Eluate was concentrated into 3C protease buffer, cleaved overnight with HRV 3C protease, and quenched with 10% (v/v) acetic acid. The acidified solution was poured over a 100 mg SEP-PAK C18 column (Waters) and washed with 3 mL 0.1% TFA in water followed by 1.5 mL each 10, 20, 30, 40, 50, 60, and 70% ACN in 0.1% TFA at room temperature. Fractions containing RPN2 peptide were identified by MALDI-TOF MS, lyophilized, and then reconstituted in water.

### Co-Purification of UCH37 and RPN13

UCH37 and RPN13 constructs were expressed in BL21(DE3)pLysS *E.coli* cells in LB media supplemented with appropriate antibiotics at 37°C to OD600 ∼0.6 and transferred to 20°C for 16 h after induction with IPTG. Cultures were harvested and stored at −80°C with UCH37•RPN13 complexes mixed 1:1 (volume) prior to lysis. Cell pellets were resuspended in lysis buffer F (50 mM HEPES pH7.5, 200 mM NaCl, 1 mM EDTA, and 1 mM TCEP), lysed by sonication, and clarified by centrifugation.

UCH37•RPN13 was purified in three chromatographic steps. (1) Clarified lysate was incubated with amylose resin for 1 h, washed with lysis buffer F, followed by low salt buffer C (50 mM HEPES pH7.5, 50 mM NaCl, and 1 mM TCEP), eluted into amylose elution buffer (low salt buffer C plus 10 mM maltose), and incubated overnight with TEV protease. (2) Eluate was incubated over Ni-NTA resin for 1 h, washed with low salt buffer C, and eluted with Ni-NTA elution buffer (lysis buffer F plus 300 mM imidazole). (3) Eluate was concentrated and run on a Superdex 200 (GE) gel filtration column in 50 mM HEPES pH7.5, 50 mM NaCl, 1 mM EDTA, and 1 mM TCEP. Minor modifications were made for the purification of isolated UCH37 and RPN13. For UCH37, step 2 was omitted and further purified using anion exchange chromatography. For RPN13, step 1 was omitted and further purified using a Superdex 75 (GE) gel filtration column.

### Purification of Proteasomes (Ptsms)

*Wild-type, UCH37^−/−^*, and *RPN13^−/−^* cells stably expressing RPN11-HTBH were grown, harvested and lysed in Ptsm buffer (40 mM HEPES pH7.4, 40 mM NaCl, 10 mM MgCl2, 2 mM ATP, 1 mM DTT, and 10% glycerol). The lysates were clarified at 20,000*xg* for 20 min and the supernatant was incubated with streptavidin resin overnight with rocking. The resin was washed and incubated for 10 min intervals with high salt wash buffer (Ptsm buffer containing 200mM NaCl) on ice with rocking. The resin was then resuspended in low salt wash buffer (Ptsm buffer without DTT) and incubated with TEV protease for 1.5 h at room temperature.

Minor modifications were made for the purification of *UCH37•RPN13-depleted, UCH37•RPN13-replenish, and UCH37 C88A•RPN13-replenish* Ptsms. For UCH37•RPN13-depleted Ptsms, clarified lysates derived from *wild-type* cells were incubated with streptavidin resin overnight in the presence of 10 µM RPN2 peptide. For UCH37•RPN13-replenish and UCH37 C88A•RPN13-replenish Ptsms, clarified lysates derived from *RPN13^−/−^* cells were incubated with streptavidin resin overnight with rocking. The resin was pelleted, resuspended in Ptsm buffer containing 10 µM WT or C88A recombinant UCH37•RPN13 complexes, and further incubated for 4 h with rocking prior to high salt washes.

To characterize purified proteasomes, western blots were used to assess individual components of the proteasome, native PAGE was used to assess intact proteasomes, Ub-AMC was used to assess deubiquitinase activity, and suc-LLVY-AMC was used to assess the chymotryptic-like peptidase activity of the proteasomes.

### Synthesis of Defined Ub chains

*Thiol-ene Trimers:* Reactions were performed as previously described (Valkevich et al., 2012). *Fluorescently labeled Native Trimers:* Reactions were performed as previously described (Crowe et al., 2016).

*Native Uniform Chains:* 2 mM Ub, 300 nM E1, 3 μM UBE2R1 (K48) or 3 μM UBE2N/UBE2V2 (K63) were mixed in reaction buffer A (20 mM ATP, 10 mM MgCl2, 40 mM Tris-HCl pH 7.5, 50 mM NaCl, and 6 mM DTT) overnight at 37°C.

*Native K6/K48 Branched Trimer:* 2mM K6/48R Ub, 1mM UbD77, 300 nM E1, 10 μM UBE2D3, and 1 μM NleL were mixed in reaction buffer A overnight at 37°C.

*Native K6/K48 Branched Tetramer*: Branched tetramer was generated using three reaction steps. (1) 2 mM K6/K48 branched tri-Ub was first cleaved by 1 µM OTUB1 in DUB buffer (50 mM Tris pH 7.5, 150 mM NaCl and 2 mM DTT) at 37°C for 1 h to generate K6 di-Ub where the proximal subunit is a UbD77 molecule. (2) 2 mM K48R Ub, 1 mM UbD77, 300 nM E1, and 3 μM UBE2R1 were mixed in reaction buffer A overnight at 37°C to generate K48 di-Ub. This K48-di Ub was then treated with 0.5 µM Yuh1 cleavage in DUB buffer at 25°C for 4 h to expose its proximal C-terminus. (3) 500 µM K6 di-Ub, 500 µM K48 di-Ub, 300 nM E1, and 3 μM UBE2R1 were mixed in reaction buffer A overnight at 37°C.

All reactions for native chains were quenched by lowering the pH to <5 via the addition of 5 M ammonium acetate pH4.4. Enzymes were then precipitated through multiple freeze thaw cycles and further purified using cation exchange chromatography.

### Generation of High Molecular Weight (HMW) Ub chains

*K6/K48 Ub chains* were assembled in a reaction buffer A containing 1mM Ub, 150 nM E1, 5 μM UBE2D3, and 3 μM NleL. *K11/48 Ub chains* were assembled in a reaction buffer B (10 mM ATP, 10 mM MgCl2, 40 mM Tris pH 8.5, 100 mM NaCl, 0.6 mM DTT, and 10% (v/v) glycerol) containing 0.6 μM Ub, 150 nM E1, and 5 μM UBE2S-UBD. 3 μM AMSH and 0.5 μM OTUD1 were added after 3 h and the mixture was left overnight at 37°C. Prior to purification, an additional bolus of AMSH and OTUD1 was added, the mixture was incubated for 3 h at 37°C, and subjected to size exclusion chromatography to isolate products with a mass >35 kDa. These HMW K11-linked chains were then added to 0.6 μM Ub, 150 nM E1, and 3 μM UBE2R1 in reaction buffer B. *K48/K63 Ub chains* were assembled using 1 mM Ub, 150 nM E1, 5 μM UBE2R1, and 5 μM UBE2N/UBE2V2 in reaction buffer A. *K11/K63 Ub chains* were assembled from 1 mM Ub, 150 nM E1, 5 μM UBE2S-UBD as described above for K11/K48 chains. Finally, *K29/K48 chains* were generated from 1 mM Ub, 150 nM E1, 2 μM UBE2D3, and 3 μM UBE3C as previously described (Michel et al., 2015).

All Ub chains were purified using size exclusion chromatography (Superdex 75) to isolate high molecular weight chains of >35kDa used in these studies.

### Purification and ubiquitination of titin I27^V15P^ 23-K-35

Titin I27^V15P^ 23-K-35 was purified as previously described (de la Peña et al., 2019). Briefly, titin I27^V15P^ 23-K-35 was expressed in BL21(DE3)pLysS *E.coli* cells in 2xYT media containing 1% glycerol and supplemented with appropriate antibiotics at 30°C to OD600 ∼1.2-1.5, induced with IPTG, and grown for an additional 5 h. Cultures were harvested, flash frozen, resuspended in lysis buffer G (60 mM HEPES pH7.5, 100 mM NaCl, 100 mM KCl, 10 mM MgCl2, 0.5 mM EDTA, 1 mg/mL lysozyme, 2 mM PMSF, 20 mM imidazole, and 10% glycerol), lysed by sonication, and clarified by centrifugation. Clarified lysate was then incubated with Ni-NTA resin for 1 h, washed with high salt buffer B, followed by low salt buffer B, and eluted with low salt buffer B containing 300 mM imidazole. Eluate was concentrated into Ulp1 protease buffer (60 mM HEPES pH7.5 and 150mM NaCl), cleaved overnight with Ulp1, and further purified using a Superdex 200 (GE) gel filtration column in 50 mM HEPES pH7.5 and 5% glycerol.

100 µM substrate was modified with 5 µM E1, 5 µM Ubc1, 20 µM Rsp5, and 2 mM Ub in labeling buffer (60 mM HEPES pH7.5, 20 mM NaCl, 20 mM KCl, 10 mM MgCl2, and 2.5% glycerol) containing 1X ATP regeneration mix for 3 h at room temperature followed by the addition of 5 μM UBE2R1 and incubation overnight at 4°C.

### Fluorescent labeling of Ubiquitinated substrates

Substrates (2 mg/mL) were fluorescently labeled using cyanine5 maleimide at pH7.2 for 2 h at room temperature and quenched with excess DTT. Free dye was separated from the substrate using a Zeba spin desalting column and buffer exchanged into labeling buffer.

### Steady-State Measurements with Defined Ub chains

Stock solutions of enzymes and ubiquitin chains were prepared in assay buffer A (50 mM HEPES pH7.5, 50 mM NaCl, and 2 mM DTT). All reactions were performed at 37°C with the exception of the K48 tetra-Ub inhibition kinetics being performed at room temperature. Each sample along with a Ub and di-Ub standard were then separated on a 15% SDS-PAGE gel and followed by SYPRO^®^ Ruby staining. Gels were visualized on a Typhoon FLA 9500 (GE) and densitometry was analyzed on the Ub standards using Image Studio™. Initial velocities of Ub and di-Ub formation were converted to concentration per minute. These values were then fit to the Michealis-Menten equation using nonlinear regression in Prism 8. Error bars represent the standard deviation of three trials for each reaction performed using UCH37 and UCH37•RPN13 complexes.

### Pulldown Assay of UBE2S-UBD-FLAG and K-GG peptides

*UBE2S-UBD-FLAG*: Anti-FLAG M2 Affinity gel (50 μL) was incubated with ubiquitinated UBE2S-UBD-FLAG (250 ng/μL) in pulldown buffer for 2 h at room temperature. The resin was washed with pulldown buffer (3x) followed by minimal buffer (50 mM HEPES pH7.4 and 150 mM NaCl, 2x). Captured UBE2S-UBD-FLAG was resuspended in minimal buffer, trypsin was then added to a 1:200 (w/w) ratio and minimal proteolysis was allowed to proceed for 2.5 h at 37°C. *K-ε-GG peptides*: Pulldown was performed according to the protocol provided by CST. Briefly, 1 mg of ubiquitinated UBE2S-UBD was resuspended in urea lysis buffer (50 mM HEPES pH8.0, 50 mM NaCl, and 8 M urea), reduced with 1.25 M DTT at 55°C for 30 min, alkylated with 1:10 (v/v) iodoacetamide for 15 min, and diluted with 20 mM HEPES pH8.0 to a final concentration of 2 M urea. Trypsin (2 μg) was added to the diluted solution and digestion was allowed to proceed overnight at 37°C. Trypsin was quenched was 10% (v/v) formic acid and peptides were purified using a SEP-PAK C18 column where peptides were washed with 2 mL 0.1% TFA in water and eluted with 50% acetonitrile with 0.1% TFA. Eluate was dried using a speed vac, resuspended in IAP buffer with added K-ε-GG antibody bead slurry (40 μL), and incubated for 2 h at 4°C with rocking. The beads were washed with IAP buffer (2x), followed by washes with water (3x), eluted with 0.15% TFA (2x), and desalted using C18 StageTips then dried with a speed vac.

All pulldown assays were done at room temperature unless otherwise indicated.

### Isothermal Titration Calorimetry (ITC) Analysis

ITC measurements were performed on a MicroCal Auto-ITC200 (Malvern) at 25°C with a setting of 20 × 2 μL injections. UCH37C88A•RPN13 and Ub chains were all buffer exchanged into dialysis buffer (50 mM HEPES pH7.4, 150 mM NaCl, and 500 μM TCEP). For UCH37•RPN13 and UCH37C88A•RPN13 measurements, the syringe contained a concentration of Ub chains at 45 μM, and the cell contained UCH37•RPN13 or UCH37C88A•RPN13 at a concentration of 3 μM. Manufactured supplied Origin software (OriginLab 7 SR4) was used to fit the data to a single-site binding model and to determine the stoichiometry (N), ΔH, ΔS, and the association constant *K*a. The dissociation constant, *K*d, was calculated from *K*a.

### Native Gel Electrophoresis

Native gel electrophoresis was performed as previously described (Elsasser et al., 2005). In brief, 2.5 μg purified proteasomes were separated on 4% acrylamide gels at 100V for 3 h at 4°C. The gel was incubated in developing buffer (50 mM HEPES pH7.4, 5 mM MgCl2, 1 mM ATP and 50 μM suc-LLVY-AMC) without agitation for 30 min at 30°C. The gel was then imaged using a UV transilluminator (Bio-Rad) with the excitation for AMC at 360nm.

### Ub-AMC and suc-LLVY-AMC Assays

UCH37, UCH37•RPN13, and Ptsms were assayed for their DUB or proteolytic activity using either Ub-AMC or suc-LLVY-AMC quenched fluorescent reporter substrates respectively. Both assays were performed in black clear bottom 96 well plates. Reactions were performed by prewarming the AMC reagent (250 nM for Ub-AMC or 50 μM suc-LLVY-AMC) dissolved in assay buffer A for UCH37 and UCH37•RPN13 and assay buffer B (40 mM HEPES pH7.4, 40 mM NaCl, 10 mM MgCl2, 2 mM ATP, and 1 mM DTT) for Ptsms at 37°C for 20 min. At this point, UCH37 and UCH37•RPN13 (20nM) or Ptsms (1 μg) in their respective assay buffer were added to the appropriate wells and hydrolysis was monitored continuously for 30 min at 37°C on a fluorescence plate reader (BioTek Synergy 2, λex = 360nm, λem = 460nm).

### Debranching and Degradation Assays

*Debranching:* Stock solutions of all DUBs and Ptsms along with HMW Ub chains (250 ng/µL) were warmed to 37°C in assay buffer A (DUBs) or assay buffer B (Ptsms). Reactions were initiated by the addition of DUB or Ptsm. Time points were taken at 1 h for DUBs and 4 h for Ptsms unless otherwise indicated and quenched with either 6x Laemmli loading buffer for immunoblotting and Ub-AQUA analysis or trypsin (1:200 w/w ratio) for middle-down mass spectrometry. For Ub-AQUA analysis, samples were separated by SDS-PAGE on 12% NuPAGE Bis-Tris gels and prepared as previously described (Ries et al., 2019). For middle-down mass spectrometry, minimal proteolysis was allowed to proceed for 2.5 h at 37°C, samples were then acidified to pH2 with acetic acid to deactivate trypsin and either (1) dialyzed into water for DUB-treated samples as previously described (Valkevich et al., 2014) or (2) separated using a Sep-pak C18 column for Ptsm-treated samples as previously described (Crowe et al., 2017).

*Degradation*: Stock solutions of Ptsms along with ubiquitinated substrates (250 ng/µL) were warmed to 37°C in assay buffer B supplemented with 1X ATP-regeneration mix. For pre-treatment of ubiquitinated substrates with UCH37•RPN13, branch points were removed for 1 h at 37°C before the addition of Ptsms. Time points were taken at 1 h unless otherwise indicated and quenched with either 6x Laemmli loading buffer for gel-based degradation monitoring or 10% (v/v) formic acid for parallel reaction monitoring. For gel-based degradation, samples were separated by SDS-PAGE on homemade 10-16% tricine gels (Schägger, 2006). Cy5 fluorescence was measured on a Typhoon FLA 9500 (GE) using a pixel density of 50 µm per pixel, while total protein staining was performed using SYPRO^®^ Ruby and imaged with a pixel density of 50 µm per pixel. For parallel reaction monitoring, corresponding light and heavy samples were mixed 1:1 (volume) and samples were desalted using C18 StageTips then dried with a speed vac. All mass spectrometry samples were resuspended in 1% formic acid.

### Flow Cytometry using GFPu-1 Reporter System

*Wild-type* and *UCH37^−/−^* GFPu-1 cells were grown in 96-well plates until cells reached ∼70% confluency. HEK293 FT cells were used as a negative control. Cells were treated with either 10 µM MG132 or 0.1% DMSO for 24 h before trypsinization and dilution in PBS with 10% FBS and 1xPen/Strep. At least 10,000 events per sample were analyzed at a flow rate of 200 µL/min on an Attune NxT Acoustic Focusing Flow Cytometer (Life Technologies) equipped with 488 nm laser (530/30 nm emission filter). Data was analyzed using FlowJo version 10.4 (FlowJo, LLC). Statistical analysis was performed using Prism 8 and are represented as the mean fluorescence in BL1-A with standard deviation of three independent experiments. \*\*\**P*<0.0005 (student’s t-test).

### Pulse-Chase Experiments with L-Azidohomoalanine (AHA) labeling

*Wild-type, UCH37^−/−^, or UCH37^−/−^* cells expressing either WT or C88S UCH37 were seeded in 6-well plates. Once cells reached ∼60% confluency, cells were washed once with PBS and incubated for 1 h in methionine free media supplemented with or without 25 µM AHA. Cells were then either harvested or chased in complete DMEM (cold chase) for 4 h. To harvest, cells were incubated in 1 mM EDTA in PBS for 5 min at 37°C under 5% CO2 before removal from plates. Cells were washed three times in PBS and resuspended in RIPA buffer (10 mM Tris-HCl pH7.5, 1 mM EDTA, 0.5% Triton X-100, 140 mM NaCl, 0.1% SDS, and 0.1% sodium deoxycholate supplemented with 1X protease inhibitor cocktail and 1X phosphatase inhibitor cocktail) before flash freezing on dry ice and storing at −80°C. Pellets were lysed via freeze-thaw cycles and sonicated before centrifugation at 20,000*xg* for 20 min at 4°C. Total protein concentration was quantified using Bradford assay and then labeled with 10 µM DBCO-Cy5 for 30 min. Labeling was quenched with 6x Laemmli loading buffer and samples were separated by SDS-PAGE on 4-20% Mini-PROTEAN gels. Cy5 fluorescence and total protein staining using SYPRO^®^ Ruby was measured on a Typhoon FLA 9500 (GE).

Minor modifications were made for *wild-type* cells treated with MG132 or *UCH37^−/−^* cells expressing either WT or C88S UCH37. For *wild-type* cells treated with MG132, cells were incubated for 4 h in methionine free media supplemented with 25 µM AHA with or without the presence of 10 µM MG132. For *UCH37^−/−^* cells expressing either WT or C88S UCH37, cells were seeded in media containing 0.1 µg/mL doxycycline for 48 h. Doxycycline remained present for lentiviral cell lines after each media change.

### Ub Middle-down Mass Spectrometry (Ub MiD MS) Analysis

Minimal tryptic fragments of HMW Ub chains were either separated using an Ultimate 3000 UHPLC (Thermo Scientific) prior to analysis using Orbitrap Fusion Tribid mass spectrometer (Thermo Scientific). For separation, the UHPLC was equipped with a MASSPrep™ Micro Desalting VanGuard Pre-Column (2.1 x 5 mm, Waters). Fragments were then separated using a linear gradient of 5% to 70% B over 18 min and 70% to 95% B over 5 min (solvent A: 0.1% formic acid (FA) in water, solvent B: 0.1% FA in ACN) using a flow rate of 10 µL/min. The resolving power of the mass analyzer on the spectrometer was set at 120000. For tandem mass spectrometry (MS/MS) using ETD, individual charge states of protein molecular ions were isolated and dissociated by ETD using a 10 ms reaction time, a 2.0e5 reagent ion target, and 10% supplemental collisionally induced dissociation (CID). All spectra were processed with in-house software (MASH Suite) using a signal-to-noise (S/N) threshold of 3 and a fit factor of 70% and then validated manually (Guner et al., 2014). Percentages correspond to the relative quantification values of the 11+ charge state for all three species: Ub1-74, 1xdiGly-Ub1-74, and 2xdiGly-Ub1-74. For measuring the relative rates of debranching for HMW chains, the relative quantification values were fit to Eq 1 using Prism 8. The reported *k*obs and error bars represent the standard deviation (SD) of two replicates for UCH37•RPN13 and varying ptsms.

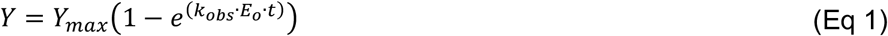

### Ubiquitin-AQUA High Resolution and Accurate Mass MS (HR/AM MS) Analysis

Full tryptic digests of HMW Ub chains were separated on an Easy nLC 1000 UHPLC equipped with a homemade 15cm nanoLC column (ProntoSIL C4 5um 300A, Custom PN-81003). Using a flow rate of 300 nL/min, the linear gradient was 0% to 50% over B for 20 min, 50% to 95% over B for 3 min, and 95% hold over B for 7 min (solvent A: 0.1% formic acid (FA) in water, solvent B: 0.1% FA in ACN). The LC system was coupled to the Orbitrap with a resolving power set at 60000. Spectra were recorded over a range 300 to 1500 m/z. For data-dependent MS/MS, the top four most intense ions with charge state of 2-5 were selected using an isolation window of 2 m/z. Fragmentation was achieved by CID at 35% nominal energy with product ion detection in the linear ion-trap. Ion chromatograms were extracted for each peptide of interest with an extraction window of 20 ppm. Chromatograms were smoothed using the Boxcar algorithm with a 7-point window. Integration was then performed using default parameters with manual adjustment as deemed appropriate. Results are normalized against total amount of Ub for each linkage type detected and are represented as means ± SEM of two replicates. For all points, asterisks represented are as follows: \**P*<0.025, \*\**P*<0.01 (student’s t-test).

### UBE2S-UBD Degradation using Parallel Reaction Monitoring (PRM) Analysis

Proteolytic peptides from UBE2S-UBD degradation were separated on an Easy nLC 1000 UHPLC equipped with a homemade 15 cm nanoLC column (ProntoSIL C4 5 um 300A, Custom PN-81003). Using a flow rate of 300 nL/min, the linear gradient was 5% to 50% over B for 35 min, 50% to 95% over B for 3 min, and 95% hold over B for 6 min (solvent A: 0.1% formic acid (FA) in water, solvent B: 0.1% FA in ACN). The LC system was coupled to the Orbitrap with a resolving power set at 50000. Spectra were recorded over a range 200 to 1300 m/z. For data-dependent MS/MS, the parent ions for the heavy (^15^N) and light (^14^N) UBE2S and UBD peptides (FPASPPKGY and FDGSGGNNHAVE) were selected using an isolation window of 2 m/z and a resolving power set at 15000. Fragmentation was achieved by HCD at a stepped 24, 27, and 30% nominal energy with transition ion detection in the orbitrap. Proteolytic peptides were first identified with an extraction window of 20 ppm using Proteome Discoverer 2.3 and ion chromatograms were then extracted each transition ion of interest with an extraction window of 10 ppm using Pinpoint 1.4. Results are represented as sum ± SD of four replicates.

## Supplemental Information

**Figure S1.**
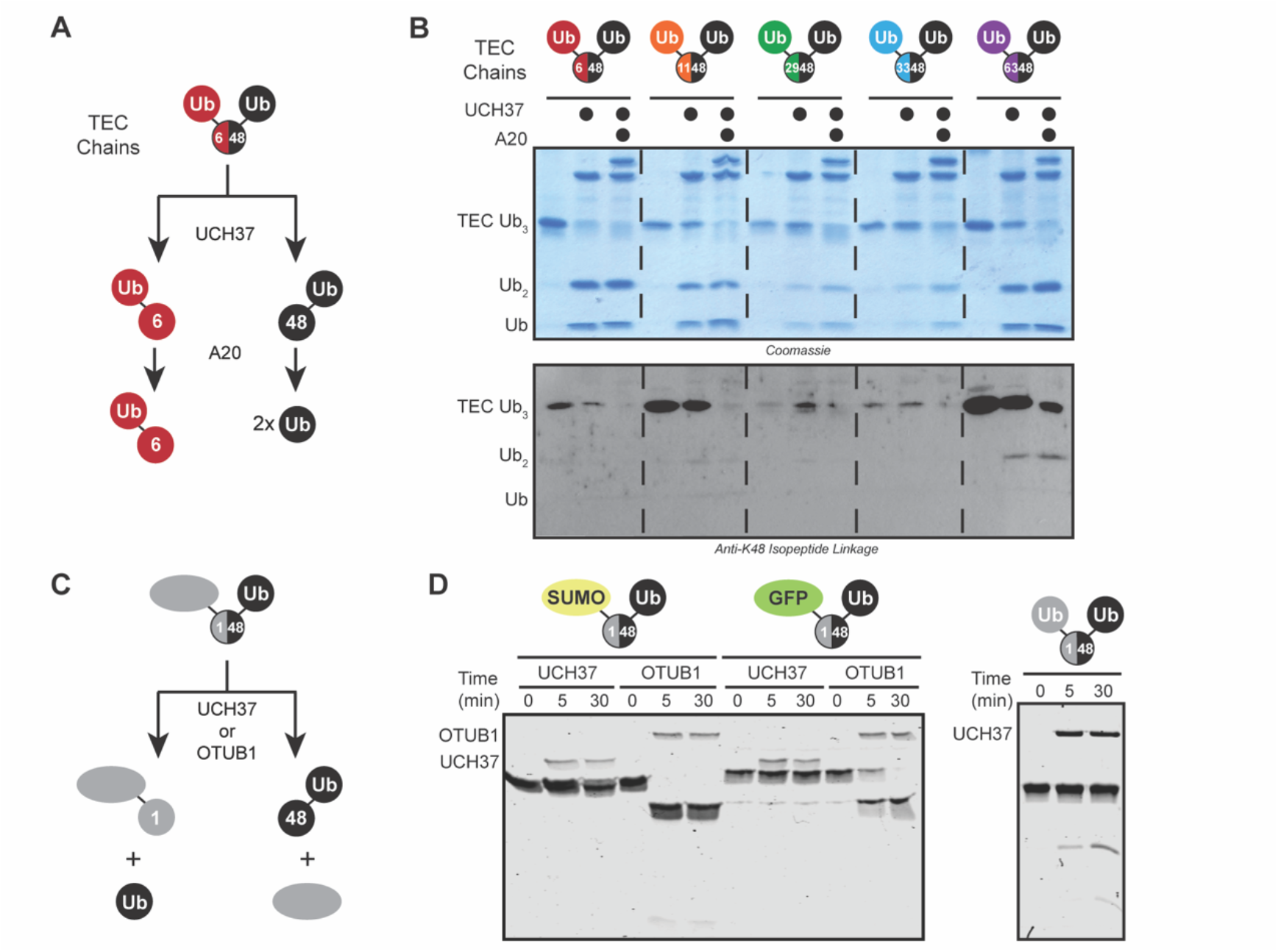
Linkage Selectivity of UCH37 with TEC Trimers, Related to Figure 1. (A) Schematic showing the sequential DUB assay used to assess the linkage selectivity of UCH37 with thiol-ene coupling (TEC)-derived Ub trimers. (B) SDS-PAGE analysis of the sequential digests of K48 containing branched tri-Ub with UCH37 (5 μM) followed by A20 (5 μM). The linkage of the di-Ub species was visualized using the anti-Ub K48-selective antibody. (C) Schematic for assessing the hydrolytic activity of UCH37 or OTUB1 with native SUMO or GFP di-Ub fusion and TEC tri-Ub. (D) SDS-PAGE analysis of native M1/K48 SUMO di-Ub and M1/K48 GFP di-Ub (10 μM, left) and TEC M1/K48 tri-Ub (10 μM, right) by UCH37 or OTUB1 (1 μM).

**Figure S2.**
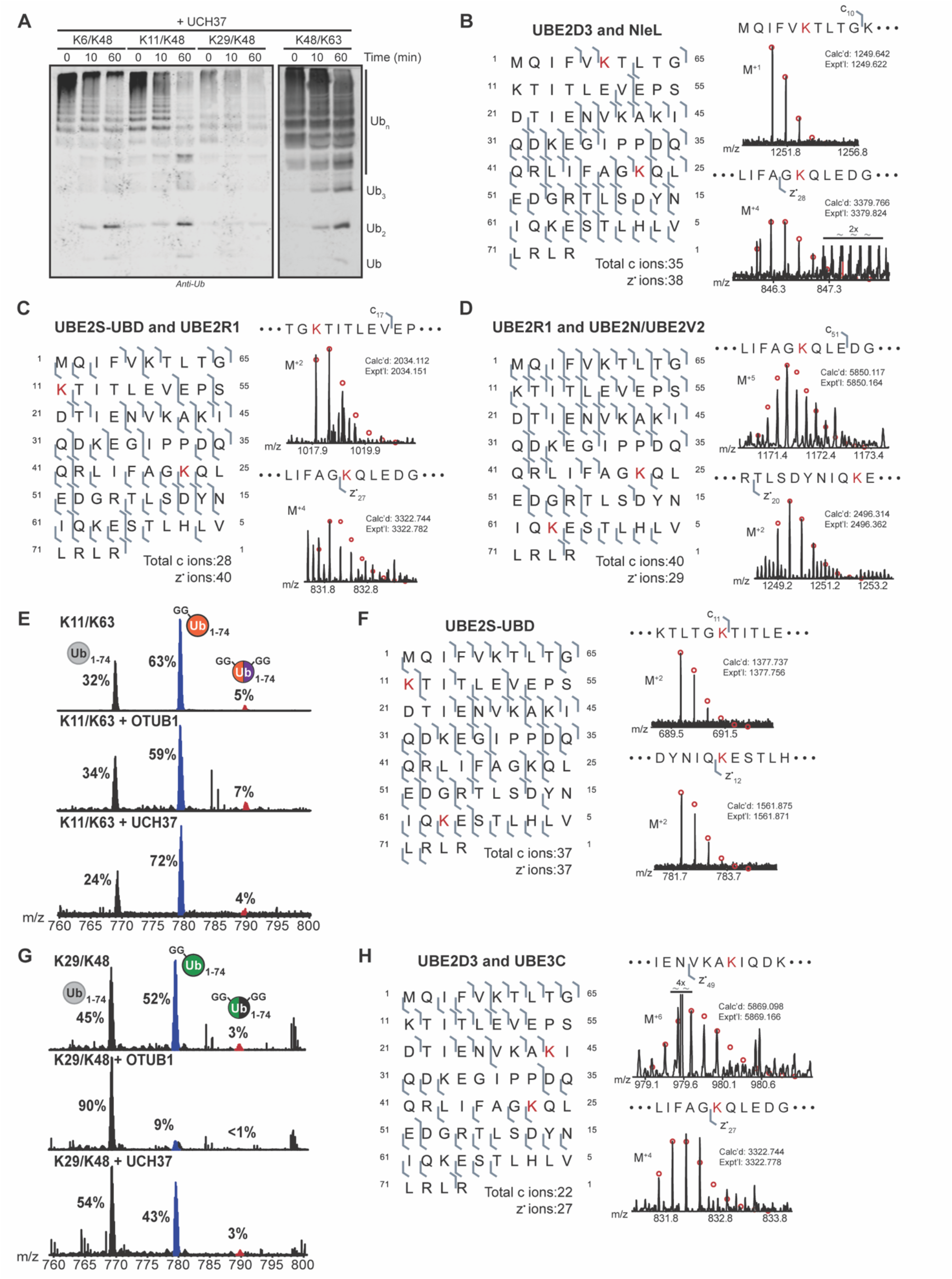
Analysis HMW Chain Cleavage with UCH37, Related Figure 2. (A) Western blot analysis of UCH37-catalyzed cleavage with HMW K6/K48, K11/K48, K29/K48, and K48/K63 Ub chains. Hydrolysis of ubiquitin chains is tracked by SDS-PAGE and visualized using the α-Ub P4D1 antibody. (B-D) Observed ETD fragments (c and z_•_ ions) mapped onto the Ub sequence containing a di-Gly modification at the following positions: K6 and K48 (B), K11 and K48 (C), and K48 and K63 (D). (E) Ub MiD MS analysis of HMW K11/K63 chains (top) treated with either OTUB1 (1 μM, middle) or UCH37 (1 μM, bottom). Percentages correspond to the relative quantification values of the 11+ charge state for each Ub species: Ub_1-74_, 1xdiGly-Ub_1-74_, and 2xdiGly-Ub_1-74_. (F) Observed ETD fragments (c and z_•_ ions) mapped onto the Ub sequence containing a di-Gly modification at K11 and K63. (G) Ub MiD MS analysis of HMW K29/K48 chains (top) treated with either OTUB1 (1 μM, middle) or UCH37 (1 μM, bottom). Percentages correspond to the relative quantification values of the 11+ charge state for each Ub species: Ub_1-74_, 1xdiGly-Ub_1-74_, and 2xdiGly-Ub_1-74_. (H) Observed ETD fragments (c and z. ions) mapped onto the sequence of Ub containing a di-Gly modification at K29 and K48. ETD fragments show the presence of a di-Gly modification (B-D, F, and H) at each respective lysine position labeled in red. Red circles represent theoretical isotopic abundance distributions of isotopomer peaks. Calc’d: calculated monoisotopic weight; expt’l: experimental monoisotopic weight.

**Figure S3.**
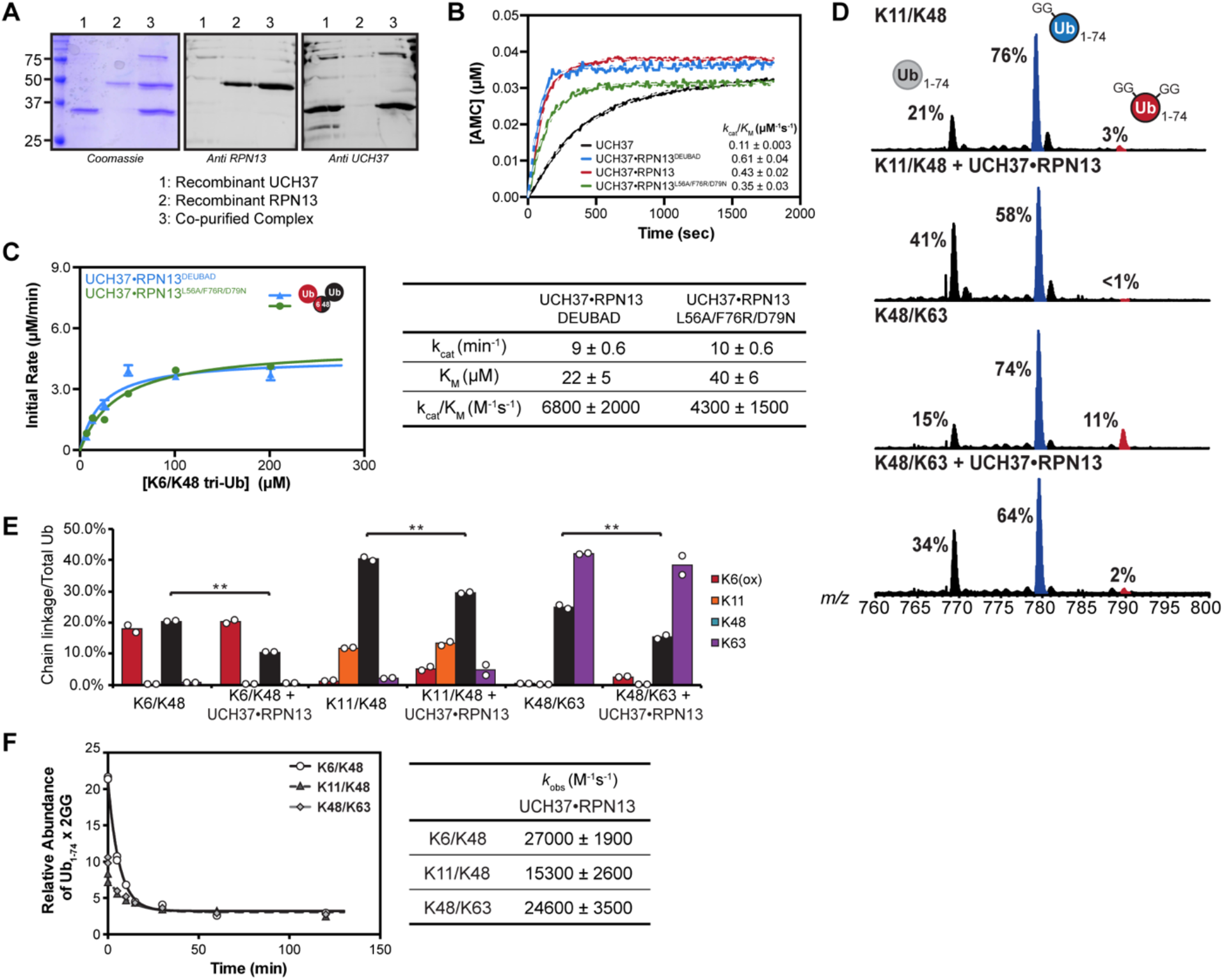
Effects of RPN13 on Debranching Activity, Related to Figure 3. (A) Characterization of the UCH37•RPN13 co-purified complex: Coomassie gel (left), α-RPN13 immunoblot (middle), and α-UCH37 immunoblot (right). 1 = recombinant UCH37, 2 = recombinant RPN13, 3 = co-purified complex. (B) Ubiquitin-AMC hydrolysis of UCH37, UCH37•RPN13^DEUBAD^, UCH37•RPN13, and UCH37•RPN13^L56A/F76R/D79N^ (20 nM). All curves are representative traces and fits are derived from averaging two independent experiments to pseudo first-order kinetics: *Y* = *_Ymax_*_(1-e(*k*cat/*K*m)•E°•t)._ (C) Michaelis-Menten plot for the hydrolysis of native K6/K48 branched tri-Ub by either UCH37•RPN13^DEUBAD^ or UCH37•RPN13^L56A/F76R/D79N^ (0.5 μM, left). Table of kinetic parameters measured for all experiments following the initial rates of di-Ub formation (right). All kinetic curves are representative traces and constants are derived from averaging fits of independent experiments with SD (n = 3). (D) Ub MiD MS analysis of HMW K11/K48 chains (top) and HMW K48/K63 chains (bottom) treated with UCH37•RPN13 (1 μM). Percentages correspond to the relative quantification values of the 11+ charge state for each Ub species: Ub_1-74_, 1xdiGly-Ub_1-74_, and 2xdiGly-Ub_1-74_. (E) Ub-AQUA analysis of HMW K6/K48, K11/K48, and K48/K63 chains before and after UCH37•RPN13 (1 μM) treatment. For all points, \**P*<0.025, \*\**P*<0.01 (Student’s T-test). Quantification values are derived from averaging fits of 2 independent experiments shown with SEM. (F) Steady-state parameters for the hydrolysis of HMW K6/K48, K11/K48, and K48/K63 chains by UCH37•RPN13 (0.5 μM, left), Catalytic efficiencies (*k*_cat_/*K*_m_) are calculated from the 11+ charge state of the 2xdiGly-Ub_1-74_ species. Table of kinetic parameters (right) measured for all experiments following the first-order decay rates of the 2xdiGly-Ub_1-74_ species. All kinetic curves are representative traces and constants are derived from averaging fits of independent experiments with SD (n = 2).

**Figure S4.**
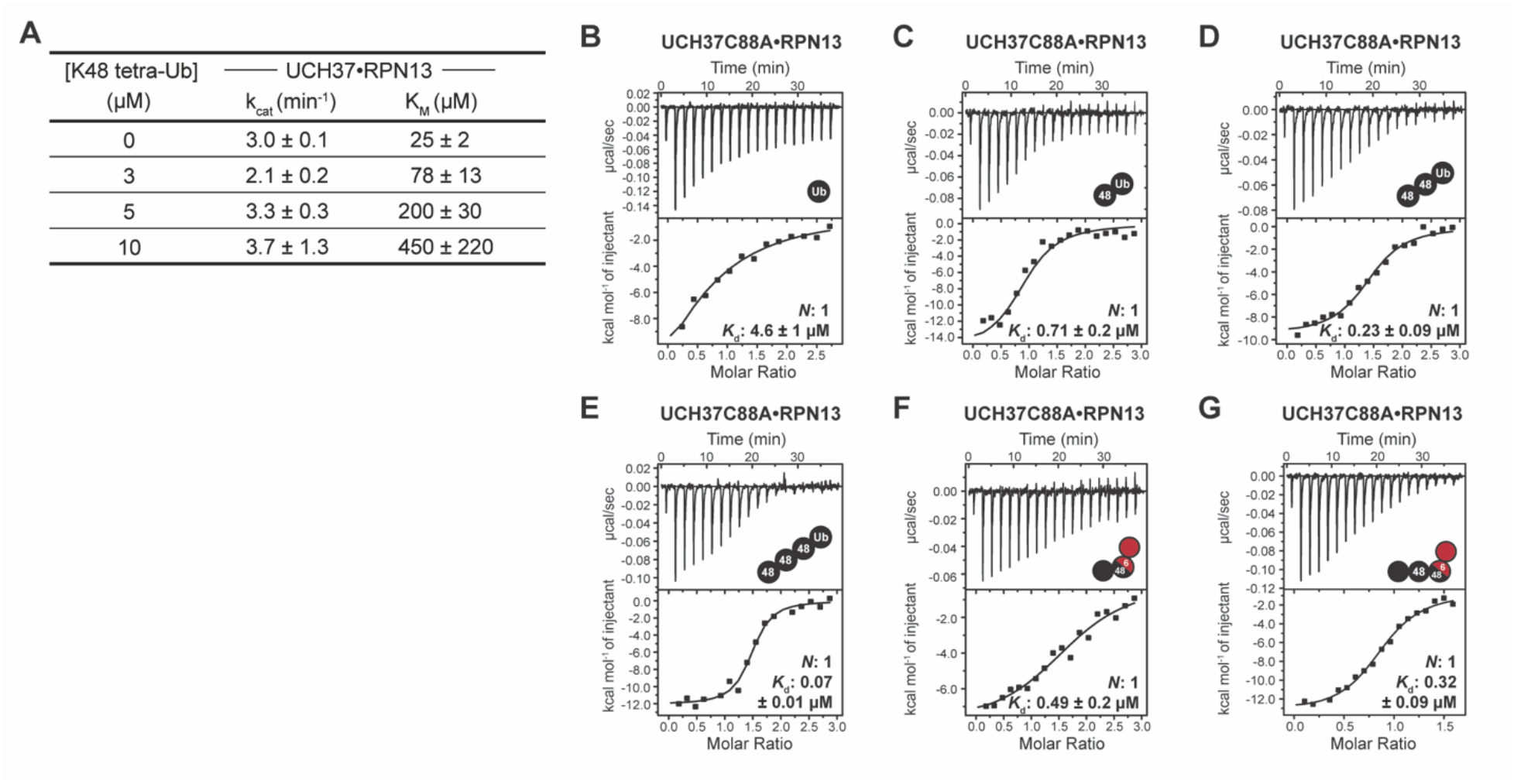
UCH37•RPN13 Binding Data, Related to Figure 4. (A) Kinetic constants derived from Michaelis-Menten analysis for the hydrolysis of native K6/K48 branched tri-Ub by UCH37•RPN13 (0.5 μM) in the presence of K48 tetra-Ub. (B-G) ITC analysis of UCH37C88A•RPN13 binding to mono-Ub (B), K48-linked di-Ub (C), tri-Ub (D), tetra-Ub (E), K6/K48 branched tri-Ub (F) and K6/K48 branched tetra-Ub (G). The reported K_d_s are derived from averaging fits of two independent experiments.

**Figure S5.**
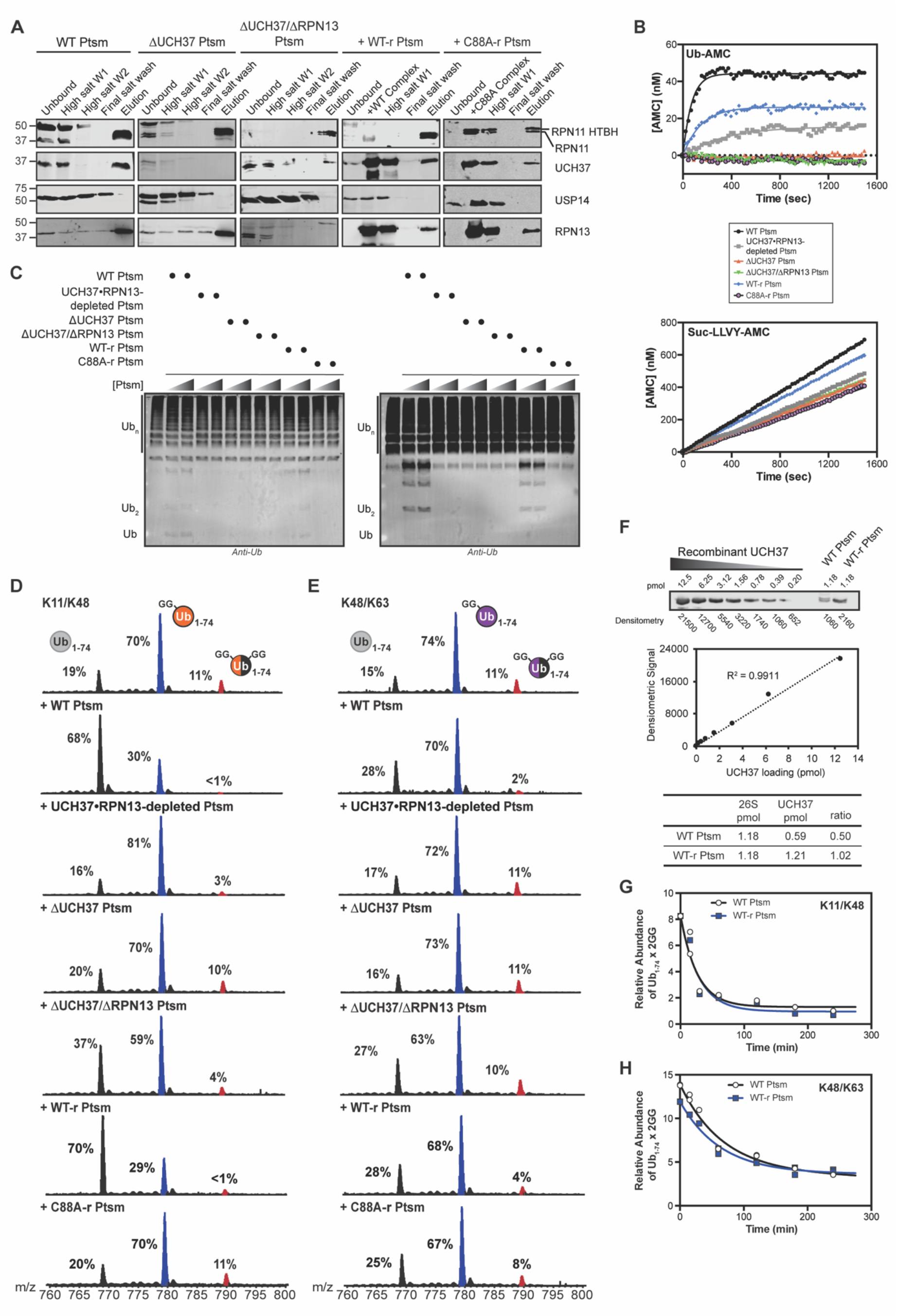
UCH37 is Required for Debranching by the Proteasome, Related to Figure 5. (A) Western blot analysis showing the loss of USP14 during purification. For KO cell lines the loss of UCH37 or RPN13 are also observed. For replenish experiments, addition of recombinant UCH37 and RPN13 are found in the final elution. (B) Ubiquitin-AMC (top) and Suc-LLVY-AMC (bottom) hydrolysis of each indicated proteasome (1 μg). All curves are representative traces and fits are derived from averaging two independent experiments. (C) Western blot analysis of HMW K11/K48 (left) and K48/K63 (right) chain debranching with increasing concentration of proteasomes using the α-Ub P4D1 antibody. (D-E) Ub MiD MS analysis of HMW K11/K48 (D) and K48/K63 (E) chains subjected to each indicated Ptsm complex (10 μg). Percentages correspond to the relative quantification values of the 11+ charge state for each Ub species: Ub_1-74_, 1xdiGly-Ub_1-74_, and 2xdiGly-Ub_1-74_. (F) Quantitative western blot analysis to determine concentration of UCH37 in WT and UCH37•RPN13-replenished proteasomes for kinetic analysis of cleavage reactions. (G-H) Steady-state parameters for the hydrolysis of HMW K11/K48 (G) and K48/K63 (H) chains by WT proteasome (10 μg, black) and UCH37•RPN13-replenished proteasome (10 μg, blue). Catalytic efficiencies (*k*_cat_/*K*_m_) are calculated from the 11+ charge state of the 2xdiGly-Ub_1-74_ species. All curves are averaged representative traces from averaging fits of independent experiments with SD (n = 2).

**Figure S6.**
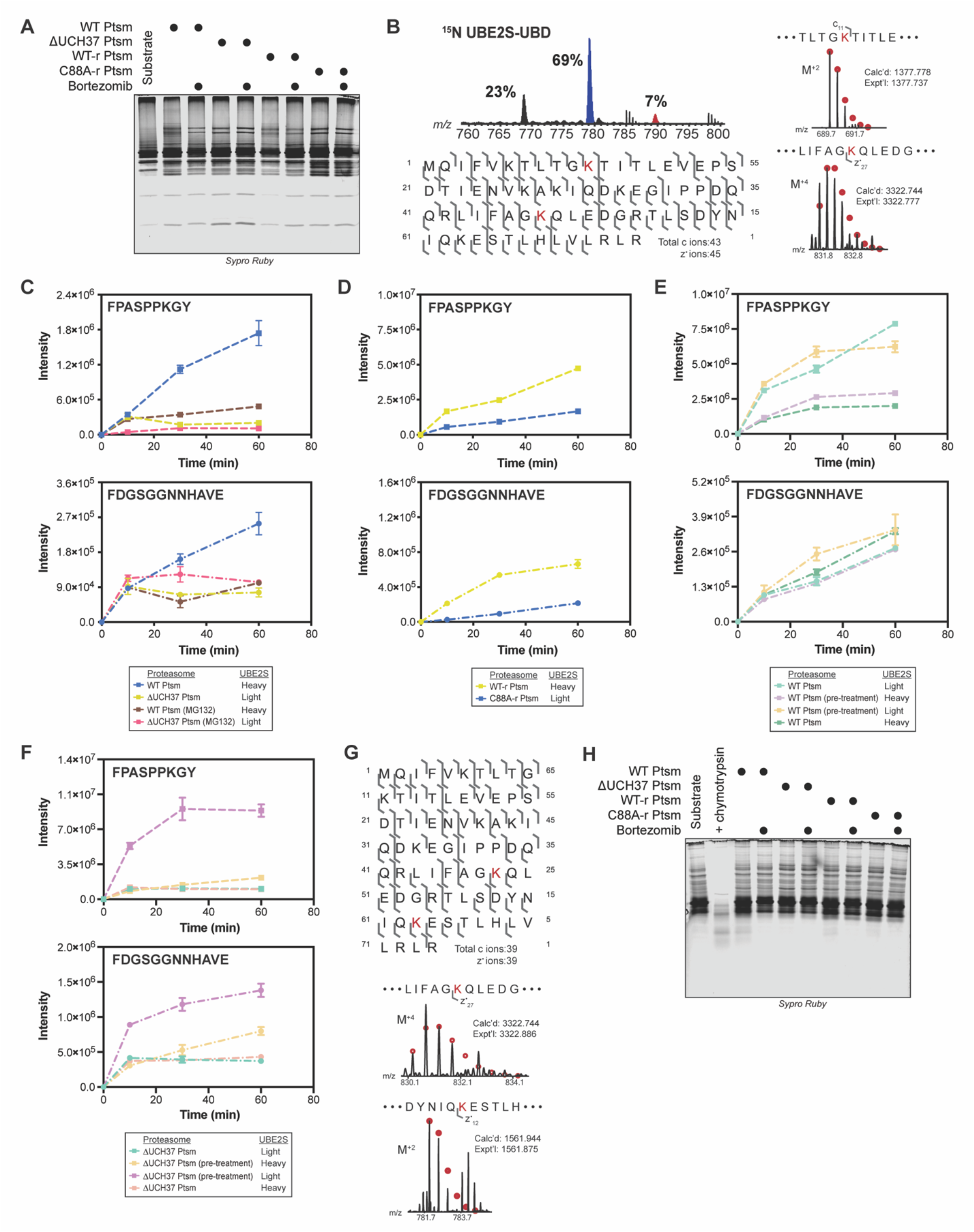
Debranching Regulates Proteasomal Degradation, Related to Figure 6. (A) Total protein stain for the degradation of K11/K48-Ub_n_-UBE2S-UBD using the indicated Ptsm complexes (5 µg) with and without the presence of bortezomib (1 μM). (B) Ub MiD MS analysis of K11/K48-Ub_n_-^15^N UBE2S-UBD (top). Percentages correspond to the relative quantification values of the 11+ charge state for each Ub species: Ub_1-74_, 1xdiGly-Ub_1-74_, and 2xdiGly-Ub_1-74_. Observed ETD fragments (c and z_•_ ions, bottom) mapped onto the Ub sequence containing a di-Gly modification at K11 and K48. (C-F) PRM MS analysis of UBE2S-UBD peptides formed by Ptsms. The UBE2S peptide FPASPPKGY is shown on the top and the UBD peptide FDGSGGNNHAVE is on the bottom. (C) Heavy and light UBE2S-UBD were mixed with either WT Ptsm (5 μg) or ΔUCH37 Ptsm (5 μg), respectively, in the presence and absence of MG132 (10 μM). (D) Light and heavy UBE2S-UBD were mixed with either WT-r Ptsm (5 μg) or C88A-r Ptsm (5 μg). (E) Light and heavy UBE2S-UBD are either untreated or pre-treated with UCH37•RPN13 (1 μM) and then mixed with WT Ptsm (5 μg). (F) Light and heavy UBE2S-UBD are either untreated or pre-treated with UCH37•RPN13 (1 μM) and then mixed with ΔUCH37 Ptsm (5 μg). (G) Observed ETD fragments (c and z_•_ ions, bottom) for the ubiquitinated titin-I27^V15P^-23-K-35 substrate mapped onto the Ub sequence containing a di-Gly modification at K48 and K63. (H) Total protein stain for the degradation of K48/K63-Ub_n_-titin-I27^V15P^-23-K-35 using the indicated Ptsm complexes (5 µg) with and without the presence of bortezomib (1 μM). ETD fragments show the presence of a di-Gly modification (B and F) at each respective lysine position labeled in red. Red circles represent theoretical isotopic abundance distributions of isotopomer peaks. Calc’d: calculated monoisotopic weight; expt’l: experimental monoisotopic weight. All MS curves are representative depictions from the sum of the transition ions of each monitored peptide (C-F) with a dashed line connecting the averaging of independent experiments with SD (n = 4). All fluorescence histograms are representative traces from averaging fits of three independent experiments.

**Figure S7.**
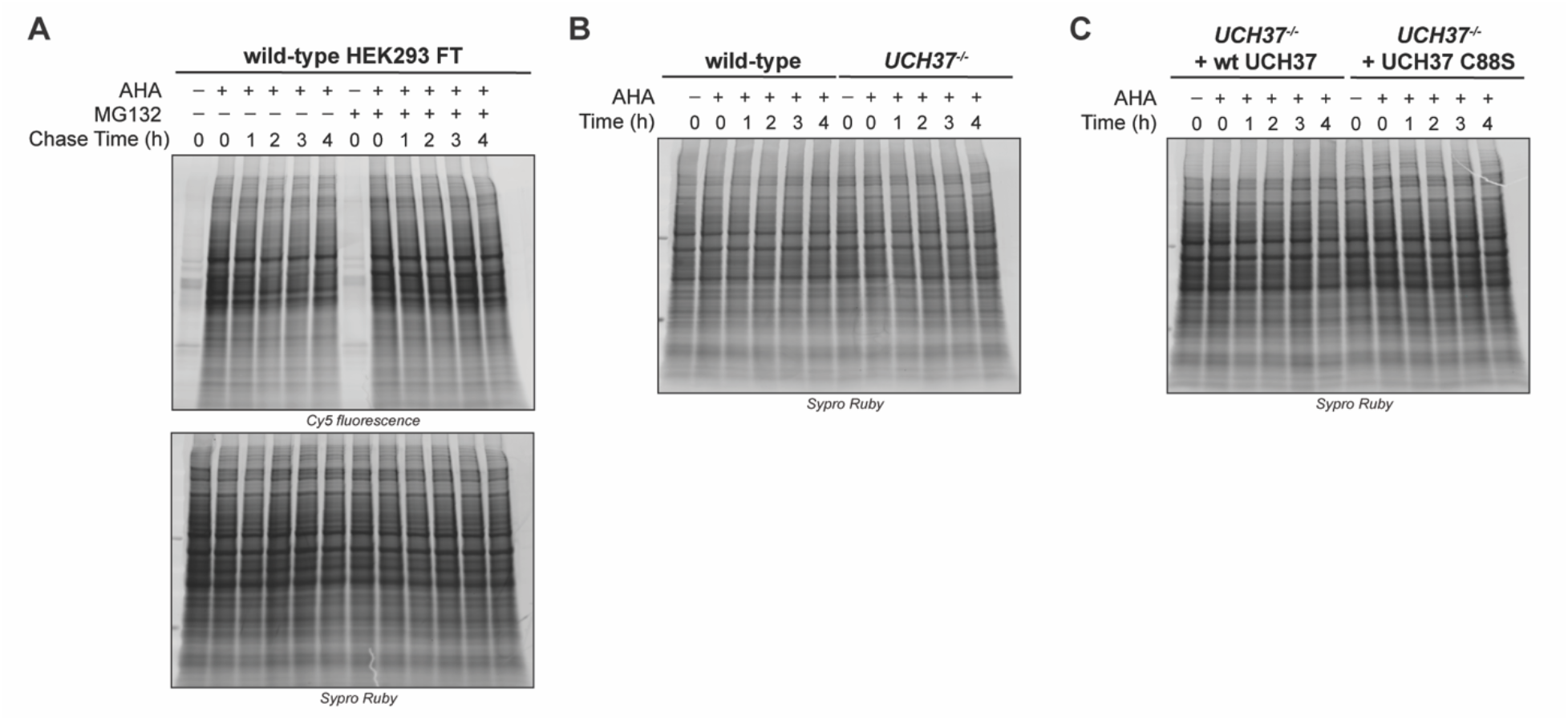
UCH37 Enhances Proteasomal Degradation, Related to Figure 7. (A) Cy5 fluorescence SDS-PAGE analysis and total protein stain of the turnover of AHA-labeled proteins at different chase times in WT HEK293 FT cells that have been treated or left untreated with MG132 (10 µM). (B) Total protein stain of the turnover of AHA-labeled proteins at different chase times in WT HEK293 FT and *UCH37^−/−^* HEK293 FT cells. (C) Total protein stain for the AHA pulse-labeling of *UCH37^−/−^* HEK293 FT cells expressing WT or C88S UCH37.

**Table S1.** Data for Ub AQUA, ITC, and PRM Analysis.

(1A) Ub peptides analyzed in this study. Peptide name, peptide sequence, accurate *m/z* of precursor ion are shown. SDS-PAGE analysis of samples analyzed in this study.

(1B) Standard curves of Ub AQUA peptides. Raw and analyzed data of ten accurate mass AQUA peptides are shown.

(1C) Ub AQUA HR/AM analysis of HMW chains analyzed in this study. Raw and analyzed data are shown for duplicate runs (related to Figure 2E-G & S3E).

(1D) ITC analysis of UCH37•RPN13 and UCH37 C88A•RPN13 complexes with K48-linked and K6/K48-linked Ub chains.

(1E) Middle-down MS analysis of HMW K6/K48, K11/K48, and K48/K63 chains with UCH37•RPN13, WT Ptsm, and UCH37•RPN13-replenish Ptsm.

(1F) MS2 analysis of K-ε-GG pulldown for K11/K48-Ub_n_-UBE2S-UBD and list of transition ions used for PRM analysis.

(1G) PRM analysis for the degradation of K11/K48-Ub_n_-UBE2S-UBD with each indicated Ptsm.

